# A NON-CANONICAL ROLE FOR NOTCH3 IN BUILDING THE INTESTINAL LYMPHATIC NICHE

**DOI:** 10.64898/2026.08.13.742813

**Authors:** Liqing Huang, Bhargav Sanketi, Madhav Mantri, Yanxi Chen, Clare Wang, Tristan Tran, Iwijn De Vlaminck, Natasza A. Kurpios

## Abstract

Lymphatic dysfunction drives severe and often intractable human diseases, yet the cellular mechanisms that establish functional lymphatic vasculature remain poorly understood. In the intestine, lacteals are specialized lymphatic vessels that absorb dietary lipids and rely on surrounding villus smooth muscle to propel lymph, forming the muscular-lacteal complex (MLC). How distinct mesenchymal populations coordinate assembly of this functional lymphatic unit remains unknown. By integrating developmental single-cell profiling, genetic lineage tracing, conditional mouse genetics, and functional assays of lipid absorption, we identify Notch3 as a central organizer of MLC development that coordinates communication between distinct mesenchymal lineages. While Notch3 promotes smooth muscle differentiation within the PDGFRα⁺ lineage, PDGFRβ⁺ lineage cells do not directly contribute to villus smooth muscle. Instead, they function as Notch3-dependent signaling hubs that instruct expansion and differentiation of neighboring PDGFRα⁺ smooth muscle progenitors via paracrine TGFβ signaling. Loss of Notch3 in PDGFRβ⁺ cells disrupts MLC development, impairs intestinal lipid absorption, and causes postnatal growth failure and lethality. Restoration of TGFβ signaling rescues the structural, functional, and survival defects caused by Notch3 loss, identifying TGFβ as a critical downstream effector of the Notch3 pathway. Furthermore, selective inhibition of canonical Notch signaling in the PDGFRβ lineage fails to phenocopy Notch3 deletion, revealing a non-canonical mechanism of Notch3 function in intestinal mesenchymal development. Together, these findings establish PDGFRβ⁺ cells as essential mesenchymal signaling organizers and define a new paradigm in which lineage-specific, non-canonical Notch3 signaling coordinates villus stromal communication to build a functional intestinal lymphatic niche.

## Introduction

Lymphatic vessels are indispensable for tissue fluid homeostasis, immune surveillance, and dietary lipid absorption. Yet despite their essential physiological functions and involvement in disorders ranging from lymphedema to obesity, diabetes, and cardiovascular disease, the lymphatic vasculature remains one of the least understood organ systems (Bernier-Latmani et al. 2015; Oliver et al. 2020; Tso et al. 2025). This knowledge gap is particularly striking in the intestine, where lacteals – the specialized lymphatic capillaries within each small intestinal villus - transport dietary lipids, lipid-soluble vitamins, lipophilic drugs, and immune cell-rich lymph into the systemic circulation (Figure 1A) (Tso and Balint 1986; Miller, McDole, and Newberry 2010).

**Figure 1.**
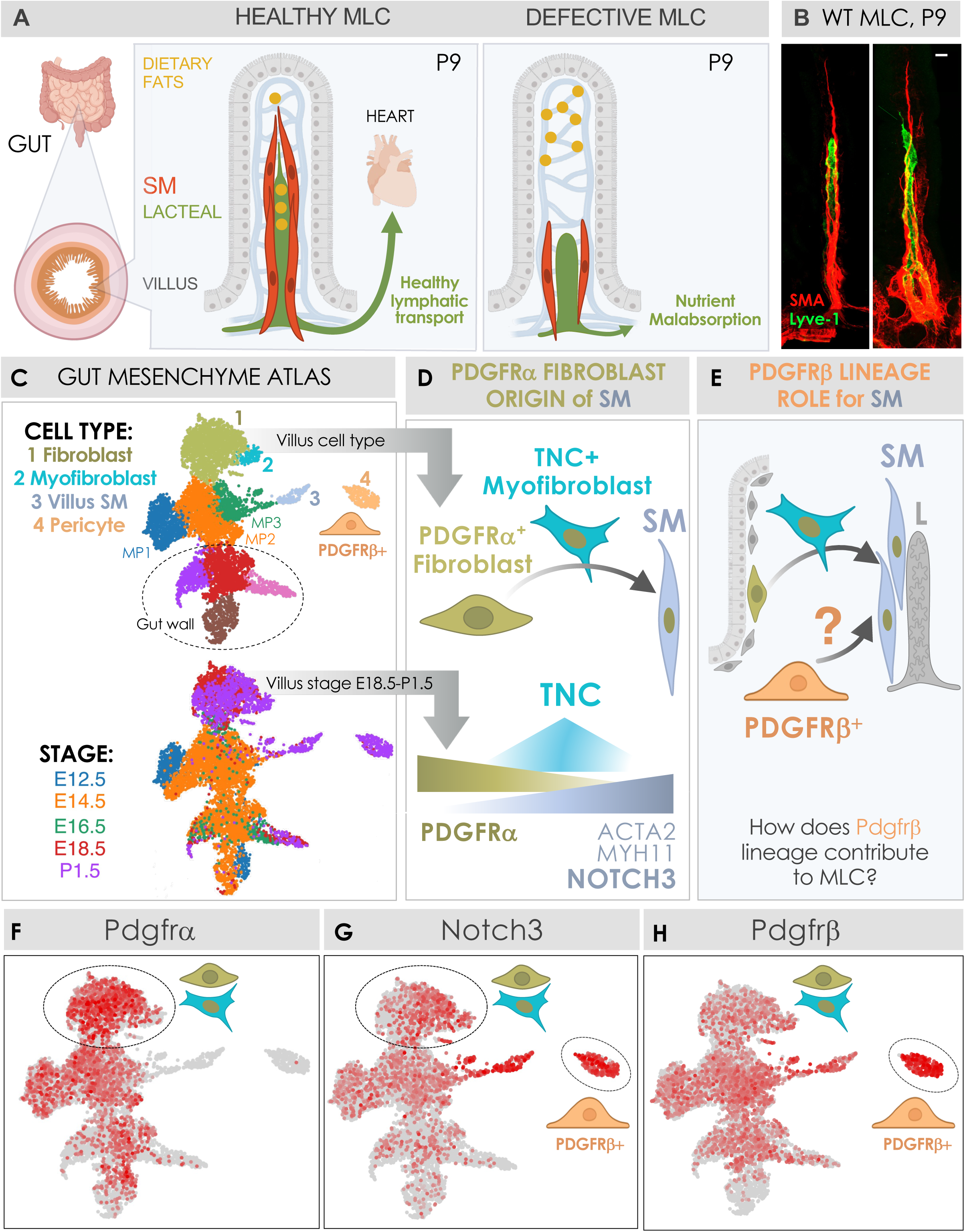
Notch3-expressing PDGFRβ lineage is associated with intestinal lymphatic niche formation. *See also Figure S1* (A) Cartoon depicting the muscular-lacteal complex (MLC) in healthy and defective postnatal day 9 (P9) intestinal villi. (B) Whole-mount immunofluorescence image of the MLC in a wild-type (WT) P9 intestine. Lacteals are labeled with LYVE-1 (green), and villus smooth muscle (SM) is labeled with αSMA (red). Arrow depicts lacteal filopodia. Scale bar, 20 μm. (C) UMAP plot of 7,519 single-cell transcriptomes from the developing mouse small intestinal mesenchyme at embryonic and postnatal days E12.5, E14.5, E16.5, and E18.5 and P1.5, clustered based on transcriptional profiles and annotated by cell identity (Sanketi et al. 2024). (D) Current model of villus SM development from PDGFRα^+^ progenitor to mature villus SM through a transient TNC⁺ myofibroblast-like intermediate “star cell” (Sanketi et al. 2024; Hu et al. 2021). (E) Additional model illustrating the potential contribution of the PDGFRβ lineage to villus SM development. (**F-H**) Feature UMAPs showing the expression of PDGFRα (**F**), Notch3 (**G**), and PDGFRβ (**H**) across the developing intestinal mesenchyme, including the pericyte cluster (marked by orange cell).

Unlike blood circulation, which is driven by the heart as a central pump, lymphatic transport depends on mechanical forces generated by surrounding contractile smooth muscle cells, making the muscular component an essential structural and functional element of lymphatic function (Scallan et al. 2016; Zawieja et al. 2025). Dysfunction of lymphatic muscle impairs lymph propulsion and contributes to lymphedema, defective immune cell trafficking, and metabolic disease (von der Weid and Zawieja 2004; Scallan et al. 2016; Petrova et al. 2004; Lutter et al. 2012; Dellinger et al. 2008; Blum et al. 2014; Davis et al. 2012). In the intestine, lymphatic transport similarly relies on the rhythmic contractions of villus smooth muscle, which mechanically propel lymph through lacteals and are required for efficient dietary lipid absorption (Figure 1A) (Tso et al. 2025; Sanders et al. 2012; Choe et al. 2015; Phan and Tso 2001; Dixon 2010; McLin, Henning, and Jamrich 2009; Thomason, Bader, and Winters 2012). We previously demonstrated that villus smooth muscle and lacteals are developmentally and functionally interdependent, forming a specialized multicellular organ-like unit - the muscular-lacteal complex (MLC) - composed of the central lacteal and its surrounding smooth muscle (Figure 1AB) (Hu et al. 2021; Sanketi et al. 2024). Disruption of this structure leads to impaired lacteal morphology with subsequent defective lipid transport and postnatal growth failure (Figure 1A) (Sanketi et al. 2024; Hu et al. 2021).

Despite the essential role of the MLC in intestinal physiology, how the intestinal mesenchyme establishes and maintains this lymphatic-supporting niche remains unclear. This knowledge gap reflects the fact that, although remarkable progress has defined epithelial diversity in the intestine, the developmental principles governing the gut mesenchyme remain comparatively unexplored. To begin addressing this question, we recently generated a developmental single-cell atlas of the intestinal mesenchyme, integrating single-cell transcriptomics, computational lineage inference, and quantitative genetic lineage tracing (Figure 1C) (Sanketi et al. 2024). These studies identified a developmental hierarchy in which the Platelet-Derived Growth Factor Receptor alpha positive (PDGFRα⁺) fibroblast-like progenitors differentiate through a transient Tenascin positive (TNC)⁺ myofibroblast-like intermediate we called “star cells” before differentiating into mature contractile villus smooth muscle (Figure 1D) (Sanketi et al. 2024; Hu et al. 2021). During this fibroblast-to-myofibroblast-to-smooth muscle transition, expression of NOTCH3, the causal gene for the hereditary vascular dementia CADASIL (Joutel et al. 1996; Kofler et al. 2015), progressively increases alongside canonical smooth muscle genes, including Acta2 and Myh11 (Figure 1D) (Sanketi et al. 2024). Consistent with this trajectory, conditional deletion of Notch3 within the PDGFRα lineage disrupts smooth muscle differentiation, prevents proper assembly of the MLC, and impairs neonatal lipid absorption (Sanketi et al. 2024). Unexpectedly, however, these developmental defects did not compromise postnatal growth or survival. Despite impaired smooth muscle differentiation and defective MLC assembly, intestinal lymphatic function was ultimately maintained (Sanketi et al. 2024). This disconnect suggested that the PDGFRα lineage does not fully account for the cellular composition or functional resilience of the MLC, pointing to previously unrecognized mesenchymal populations that contribute to its development and function. Therefore, we sought to define the additional mesenchymal programs that coordinate MLC development and support intestinal lymphatic transport.

Our developmental single-cell atlas identified a distinct mesenchymal population with features suggesting a potential role in MLC (Figure 1E). Specifically, we detected robust Notch3 expression within a Platelet-Derived Growth Factor Receptor beta positive (PDGFRβ⁺ PDGFRα⁻) cell cluster (Figure 1F-H), implicating PDGFRβ⁺ cells as potential regulators of MLC function. PDGFRβ is widely used as a marker of pericytes which, together with vascular smooth muscle cells, comprise the mural cell family (Armulik, Genové, and Betsholtz 2011; Cappellari and Cossu 2013; Majesky et al. 2011) and in select developmental contexts can contribute to smooth muscle formation in response to Notch signaling (Volz et al. 2015). Whether villus PDGFRβ⁺ cells similarly serve as direct sources of smooth muscle progenitors or instead regulate MLC development indirectly through Notch3-dependent paracrine signaling to neighboring PDGFRα⁺ progenitors remains unknown.

Here, we identify Notch3 as a central regulator of villus MLC development that performs distinct PDGFR lineage-specific functions within the intestinal mesenchyme. While Notch3 promotes smooth muscle differentiation within PDGFRα⁺ lineage, PDGFRβ⁺ cells do not serve as smooth muscle progenitors. Instead, Notch3 establishes a paracrine Transforming Growth Factor Receptor beta (TGFβ) signaling program in PDGFRβ⁺ cells that expands the PDGFRα⁺ smooth muscle progenitor population, enabling MLC assembly, dietary lipid absorption, and postnatal growth. These findings define PDGFRβ⁺ cells as critical signaling hubs that regulate villus smooth muscle development through a non-cell-autonomous mechanism and reveal a previously unrecognized, non-canonical function of Notch3 in the intestinal mesenchyme. Together, our study establishes a new paradigm in which communication between distinct mesenchymal lineages builds the intestinal lymphatic niche and provides a framework for understanding how Notch3-dependent stromal signaling may contribute to vascular remodeling and smooth muscle dysfunction in human disease.

## Results

### A Notch3-expressing PDGFRβ lineage is associated with intestinal lymphatic niche formation

The mechanisms that coordinate MLC development remain incompletely understood. Although we previously demonstrated that Notch3 is required within the PDGFRα lineage for villus smooth muscle differentiation and MLC assembly, deletion of Notch3 in these cells disrupted dietary lipid absorption without impairing postnatal growth. This finding suggested that additional Notch3-dependent mechanisms cooperate with the PDGFRα lineage to establish a functional MLC. Our developmental single-cell atlas revealed robust Notch3 expression within the PDGFRβ lineage throughout intestinal development (Figure 1GH). Although PDGFRβ is widely used as a marker of pericytes (Figure S1), its expression is dynamically regulated during intestinal development. PDGFRβ broadly marks mesenchymal populations during early embryogenesis (E12.5–E14.5; Figure 1C and H), as well as PDGFRα⁺ fibroblast and myofibroblast-like intermediates at later stages (E16.5–P1.5; Figure 1C and H), before becoming progressively enriched in PDGFRα⁻NG2⁺ pericytes during early postnatal development (PDGFRβ^high^, P1.5; Figure 1C and H; Figure S1). This pattern suggested that Notch3 functions in both PDGFRα- and PDGFRβ-derived mesenchymal lineages to coordinate establishment of the intestinal lymphatic niche (Figure 1E).

### Loss of Notch3 in the PDGFRβ lineage causes postnatal growth failure and lethality

To investigate the role of Notch3 in the PDGFRβ lineage, we conditionally deleted Notch3 (Figure 2A) using the PDGFRβ-CreERT2 driver (Notch3ΔPdgfrβ) following the same tamoxifen induction-collection interval (P0i-P9c) as previously used for PDGFRα-specific deletion (Notch3ΔPdgfrα) (Sanketi et al. 2024). We monitored body weight throughout early postnatal development to determine the impact of Notch3 loss on growth (Figure 2B). No significant differences were detected at P4 (Control: 2.123 ± 0.03 g, n = 26; Notch3ΔPdgfrβ: 2.05 ± 0.04 g, n = 26; Notch3ΔPdgfrα: 2.23 ± 0.04 g, n = 26). However, by P9, Notch3ΔPdgfrβ pups displayed a pronounced reduction in body weight compared with littermate controls (Control: 4.163 ± 0.08 g, n = 26; Notch3ΔPdgfrβ: 2.96 ± 0.10 g, n = 26), whereas Notch3ΔPdgfrα mice remained indistinguishable from controls (3.98 ± 0.14 g, n = 26) (Figure 2BC). Consistent with the increased severity of the growth phenotype over Notch3ΔPdgfrα mice, Notch3ΔPdgfrβ mice experienced a 70% mortality rate at P30 (Figure S2), demonstrating that Notch3 signaling in the PDGFRβ lineage is essential not only for normal postnatal growth but also for postnatal survival.

**Figure 2.**
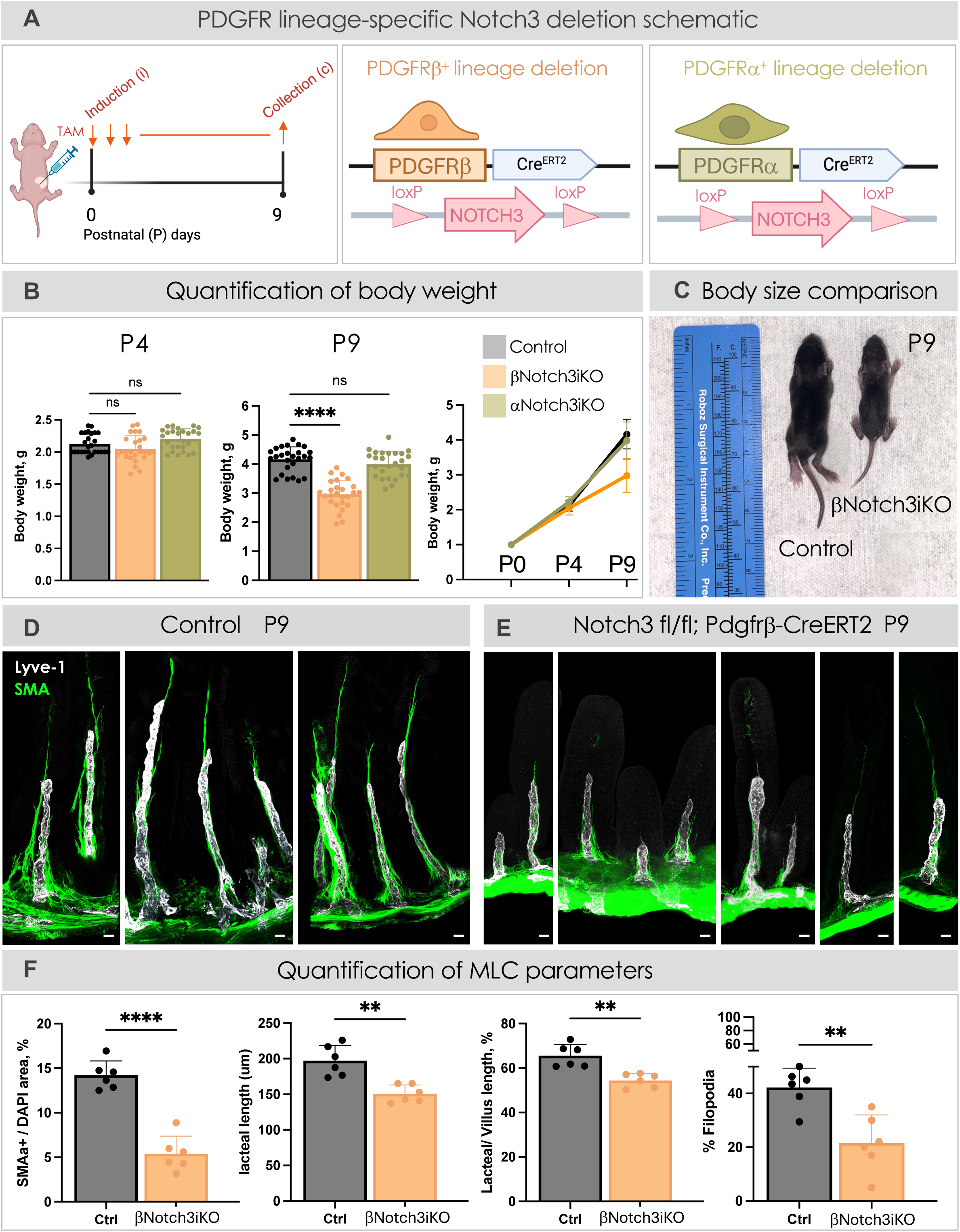
PDGFRβ-lineage Notch3 loss disrupts MLC, growth, and survival. *See also Figure S2* (A) Schematic illustrating the generation of inducible Notch3ΔPdgfrβ and Notch3ΔPdgfrα mice. PDGFRβ^+^- and PDGFRα^+^ cell-specific depletion of Notch3 starting at P0 and tissue collection at P9 (P0-P9). (B) Distinct effects of Notch3 deletion in PDGFRβ⁺ and PDGFRα⁺ lineages on postnatal growth. Body weights of control, Notch3ΔPdgfrβ (βNotch3iKO), and Notch3ΔPdgfrα (αNotch3iKO) mice were measured at P4 and P9. <u>P4</u>: No significant differences in body weight were observed among the three groups. P4 control: 2.123 ± 0.03 g; P4 Notch3ΔPdgfrβ: 2.05 ± 0.04 g; P4 Notch3ΔPdgfrα: 2.23 ± 0.04 g, p= 0.0104. <u>P9:</u> Notch3ΔPdgfrβ pups exhibited a significant reduction in body weight compared with control littermates, P9 control:4.163 ± 0.08 g; P9 Notch3ΔPdgfrβ: 2.96 ± 0.10 g, ****p< 0.0001, whereas Notch3ΔPdgfrα mice showed no significant change: 3.98 ± 0.14 g, p=0.3156 (ns). The growth curves (right) illustrate comparable body weights among all groups at birth (P0) and P4, followed by impaired postnatal weight gain specifically in Notch3ΔPdgfrβ mice between P4 and P9. Dots indicate biological replicates (n=26), pooled from six independent experiments. Data are presented as mean ± SEM. Statistical significance was determined by one-way ANOVA followed by Dunnett’s multiple-comparisons test. ns, not significant. Control genotype: PDGFRβ-Cre/+; Rosa26-tdTomato. (C) A pair of pups showing significant growth retardation of Notch3ΔPdgfrβ versus the control at P9. (**D, E**) Whole-mount MLC of Control (**D**) and Notch3ΔPdgfrβ (**E**) at P9. White is lacteal (Lyve-1); Green is αSMA (smooth muscle). Scale bar, 20 μm. Control genotype: PDGFRβ-Cre/+; Rosa26-tdTomato. (F) Comparisons of the MLC parameters in control and Notch3ΔPdgfrβ mice at P9. Villus SM area per villus: Control: 14.21% ± 1.61%; Notch3ΔPdgfrβ: 5.40% ± 1.97%, ****p <0.0001. Lacteal length: Control: 197.2 ± 8.77; Notch3ΔPdgfrβ: 150.7 ± 5.03, **p=0.0018. Lacteal length normalized to villus length: Control: 65.55% ± 2.08%; Notch3ΔPdgfrβ: 54.46% ± 1.27%, **p = 0.0017. Percent lacteals with filopodia in the jejunum: Control: 42.19% ± 2.95%; Notch3ΔPdgfrβ: 21.50% ± 4.28%, **p = 0.0033. Dots indicate values from 90-120 villi/group from three independent experiments using n=6 mice/group. Data are presented as mean ± SEM. Statistical significance was determined by unpaired two-tailed Student’s *t*-tests with Welch’s correction. Control genotype: PDGFRβ-Cre/+; Rosa26-tdTomato.

### Loss of Notch3 in the PDGFRβ lineage disrupts muscular-lacteal complex development

Because postnatal growth depends on MLC (Hu et al. 2021; Sanketi et al. 2024), we first examined villus smooth muscle, a critical structural component of this complex. Compared with control littermates, Notch3ΔPdgfrβ mice exhibited a marked reduction in villus smooth muscle coverage (Figure 2D-F; Control: 14.21% ± 1.61%, n = 6; Notch3ΔPdgfrβ: 5.40% ± 1.97%, n = 6), indicating that Notch3 in the PDGFRβ lineage is required for normal smooth muscle development. We next asked whether this smooth muscle defect was accompanied by abnormalities in the associated lacteals. In control mice, lymphatic endothelial cells (LECs) at the lacteal tip extended abundant filopodia, specialized protrusions required for lacteal elongation and regeneration (Hu et al. 2021; Bernier-Latmani et al. 2015). In contrast, lacteals in Notch3ΔPdgfrβ mice displayed a marked reduction in filopodia number at P9 (Figure 2D-F; Control: 42.19% ± 2.95%, n = 6; Notch3ΔPdgfrβ: 21.50% ± 4.28%, n = 6). Consistent with impaired lacteal growth, mutant lacteals were significantly shorter than those of control littermates (Figure 2D-F; Lacteal length: Control: 197.2 ± 8.77, n = 6; Notch3ΔPdgfrβ: 150.7 ± 5.03, n = 6; Normalized lacteal length: Control: 65.55% ± 2.08%, n = 6; Notch3ΔPdgfrβ: 54.46% ± 1.27%, n = 6). Together, these findings demonstrate that Notch3 signaling in the PDGFRβ lineage is required for coordinated development of villus smooth muscle and lacteals, thereby preserving the structural integrity of the MLC.

### Loss of Notch3 in the PDGFRβ lineage disrupts lacteal junctional specialization

Lacteals serve as the primary route for dietary lipid absorption, a function that depends on specialized LEC junctional architecture (Baluk and McDonald 2022). To determine whether Notch3 deletion in the PDGFRβ lineage altered lacteal function, we examined VE-cadherin localization in jejunal villi and classified LEC junctions as button, intermediate, or zipper structures based on their morphology (Zarkada et al. 2023; Jannaway et al. 2023). Loss of Notch3 in the PDGFRβ lineage significantly altered lacteal junctional organization, shifting junctions away from the highly permeable button configuration toward less permissive intermediate and zipper states (Figure 3A-D). Specifically, Notch3ΔPdgfrβ lacteals exhibited a significant reduction in button junctions (Control: 84.33% ± 1.93%, n = 30; Notch3ΔPdgfrβ: 52.88% ± 3.25%, n = 59), accompanied by increased intermediate junctions (Control: 14.00% ± 1.60%, n = 30; Notch3ΔPdgfrβ: 44.92% ± 1.88%, n = 59) and zipper junctions (Control: 1.17% ± 0.52%, n = 30; Notch3ΔPdgfrβ: 8.81% ± 1.84%, n = 59). In contrast, deletion of Notch3 in the PDGFRα lineage did not alter the distribution of lacteal junctional subtypes (Figure 3A-D), demonstrating a lineage-specific requirement for PDGFRβ-associated Notch3 in regulating lacteal endothelial specialization. Together, these findings reveal that PDGFRβ-lineage Notch3 is required for establishment of the highly permeable lacteal junctional architecture that supports efficient dietary lipid uptake.

**Figure 3.**
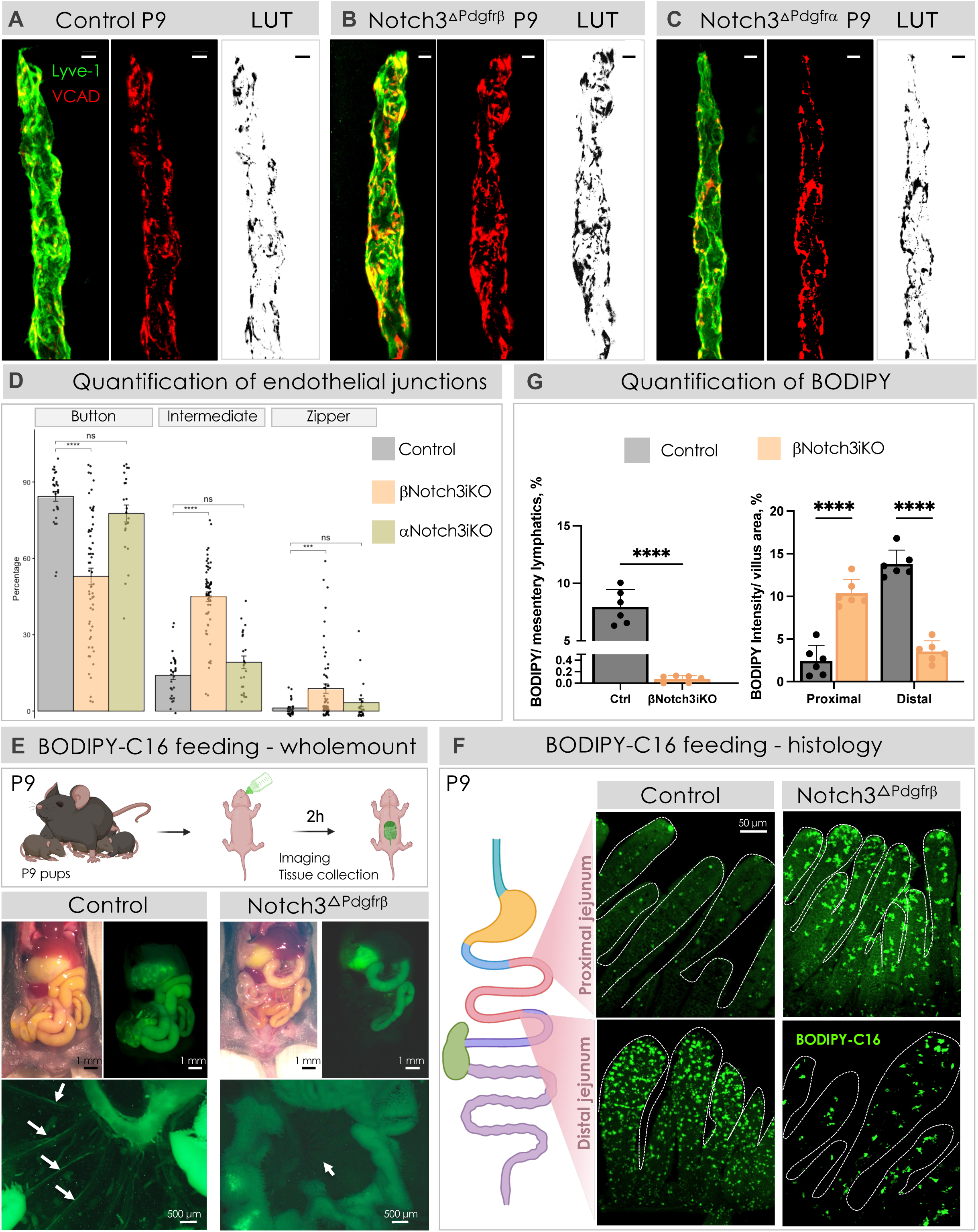
PDGFRβ-lineage Notch3 is required for dietary lipid absorption. *See also Figure S3* (**A-C**) Representative images of VE-cadherin+ (VCAD, red) lymphatic endothelial cell (LEC) junctions of LYVE-1+ lacteals (green) in control (**A**), Notch3ΔPdgfrβ (**B**), and Notch3ΔPdgfrα (**C**) mice. Scale bars, 5 μm. Control genotype: Notch3fl/fl. (D) Quantification of % button, intermediate, and zipper-like junctions out of total junction length in lacteals shown in A-C. Each dot represents one lacteal (control: 30 lacteals; Notch3ΔPdgfrβ: 59 lacteals; Notch3ΔPdgfrα: 23 lacteals, 6 mice per group). <u>Button</u>: control: 84.33% ± 1.93%; Notch3ΔPdgfrβ: 52.88% ± 3.25%; Notch3ΔPdgfrα: 77.61% ± 3.34%. <u>Intermediate</u>: control: 14.00% ± 1.60%; Notch3ΔPdgfrβ:44.92% ± 1.88%; Notch3ΔPdgfrα: 19.13% ± 2.49%. <u>Zipper</u>: control: 1.17% ± 0.52%; Notch3ΔPdgfrβ: 8.81% ± 1.84%; Notch3ΔPdgfrα: 3.26% ± 1.56%. Group differences in junction composition were analyzed by one-way ANOVA, followed by pre-specified pairwise Student’s *t*-tests comparing control with Notch3ΔPdgfrβ (βNotch3iKO), and control with Notch3ΔPdgfrα (αNotch3iKO). For junction-length measurements in which multiple observations were obtained from each mouse, data were analyzed using a linear mixed-effects model with mouse as a random effect, and *P* values were calculated using Satterthwaite’s method. Data are presented as mean ± SEM. Statistical significance was determined by unpaired t-test. *** p = 0.000164, **** p <0.0001. ns, not significant. Control genotype: Notch3fl/fl. (E) Diagram depicting the schedule of BODIPY C16 in P9 control or Notch3ΔPdgfrβ mice (top). Fluorescent imaging of lipid tracer BODIPY C16 in the abdominal cavity (<u>middle</u>; Scale bar, 1 mm) and mesenteries (<u>bottom</u>; Scale bar, 500 μm). Note the presence of the tracer in the distal ileum in the control versus in the mid-jejunum in Notch3ΔPdgfrβ. (G, <u>left</u>) Quantification of tracer-filled lymphatics in mesenteries of control and Notch3ΔPdgfrβ mice. Control: 7.94% ± 0.61%; Notch3ΔPdgfrβ: 0.075% ± 0.024%. Each dot represents one mouse, n = 6 per group, mice pooled from two independent experiments. Data are presented as mean ± SEM. Statistical significance was determined by unpaired two-tailed Student’s t-tests with Welch’s correction. **** p <0.0001. Control genotype: Notch3fl/fl. (F) Fluorescent imaging of BODIPY C16 in the fixed proximal (top) and distal (bottom) jejunal villi in P9 control or Notch3ΔPdgfrβ. BODIPY C16 was found to accumulate in the proximal jejunum villi of Notch3ΔPdgfrβ, while the signal is concentrated in the distal jejunum of control. (G, right) Quantification of BODIPY C16 signal in the villi of control and Notch3ΔPdgfrβ mice. <u>Proximal jejunum</u>: Control: 2.45% ± 0.74%; Notch3ΔPdgfrβ: 10.37% ± 0.66%; <u>Distal jejunum</u>: Control: 13.8% ± 0.66%; Notch3ΔPdgfrβ: 3.54% ± 0.51%. Each dot represents one mouse, n = 6 per group, mice pooled from two independent experiments. Data are presented as mean ± SEM. Statistical significance was determined by unpaired two-tailed Student’s t-tests with Welch’s correction. **** p <0.0001. Scale bars, 50 μm. Control genotype: Notch3fl/fl.

### Notch3 in the PDGFRβ lineage is required for dietary lipid absorption

The altered lacteal junctional architecture suggested that loss of Notch3 in the PDGFRβ lineage may compromise the primary function of lacteals. We therefore examined whether disruption of MLC in Notch3ΔPdgfrβ mice resulted in measurable defects in lipid absorption. First, we orally administered fluorescent BODIPY C16 to neonatal mice (Figure 3E). Two hours after administration, BODIPY C16 had reached the distal ileum in control pups but was detected only in the mid-jejunum of mutant littermates, indicating delayed intestinal lipid transport (Figure 3E). Furthermore, BODIPY fluorescence was nearly absent from the mesentery of mutant mice (Figure 3E and G; Control: 7.94% ± 0.61%, n = 6; Notch3ΔPdgfrβ: 0.075% ± 0.024%, n = 6), indicating markedly impaired lymphatic lipid uptake and transport. Whole-mount imaging of villi further demonstrated that BODIPY fluorescence was predominantly localized to the distal jejunum in control animals, whereas in mutant mice it remained confined to the proximal jejunum (Figure 3FG; Proximal jejunum: Control: 2.45% ± 0.74%, n = 6; Notch3ΔPdgfrβ: 10.37% ± 0.66%, n = 6; Distal jejunum: Control: 13.8% ± 0.66%, n = 6; Notch3ΔPdgfrβ: 3.54% ± 0.51%, n = 6), further supporting delayed intestinal lipid transit. Collectively, these results demonstrate that Notch3 deletion in the PDGFRβ lineage impairs both lacteal permeability and lymphatic lipid transport, resulting in defective dietary lipid absorption.

### Postprandial serum lipid profiling confirms impaired dietary lipid absorption in the absence of PDGFRβ-lineage Notch3

To further validate the defect in dietary lipid absorption, pups were challenged with an oral lipid gavage followed by analysis of postprandial serum lipid levels (Figure S3). Compared with control littermates, Notch3ΔPdgfrβ mice exhibited significantly reduced serum triglyceride levels following lipid challenge (Control: 409.7 ± 41.4 mg/dL, n = 4; Notch3ΔPdgfrβ: 256.8 ± 19.07 mg/dL, n = 4), consistent with impaired intestinal lipid uptake. In addition, Notch3ΔPdgfrβ mice displayed altered postprandial serum lipid profiles, including changes in total cholesterol (Control: 89.5 ± 1.56 mg/dL, n = 4; Notch3ΔPdgfrβ: 114.3 ± 4.91 mg/dL, n = 4), HDL (Control: 27.25 ± 0.85 mg/dL, n = 4; Notch3ΔPdgfrβ: 27.75 ± 0.75 mg/dL, n = 4), and LDL levels (Control: 15.75 ± 1.03 mg/dL, n = 4; Notch3ΔPdgfrβ: 21.0 ± 1.47 mg/dL, n = 4). Together, these findings independently validate that loss of Notch3 in the PDGFRβ lineage disrupts intestinal dietary lipid absorption and systemic lipid handling.

### Intestinal PDGFRβ lineage is not the major source of villus smooth muscle progenitors

The disruption of villus smooth muscle following Notch3 deletion in the PDGFRβ lineage raised an important mechanistic question: do PDGFRβ⁺ mesenchymal cells directly contribute to villus smooth muscle formation, or do they regulate smooth muscle development through a non-cell-autonomous mechanism (Figure 4A)? Our previous studies established that villus smooth muscle arises from the PDGFRα lineage through a developmental transition in which PDGFRα⁺ fibroblast-like progenitors differentiate into TNC⁺ myofibroblast-like star cells before maturing into contractile villus smooth muscle cells (Sanketi et al. 2024). To determine whether the PDGFRβ lineage similarly contributes to villus smooth muscle formation, we performed lineage tracing using PDGFRβ-CreERT2; Rosa26-tdTomato reporter mice by intraperitoneal administration of tamoxifen (TAM) in neonatal mice and during adulthood. As an essential validation step, we first performed short-term lineage tracing by intraperitoneal administration of 4-hydroxytamoxifen (4OHT; the active metabolite of tamoxifen) to define the initial PDGFRβ-expressing population and establish the specificity of our labeling strategy. Short-term labeling of PDGFRβ-CreERT2; Rosa26-tdTomato mice in neonatal and adult mice revealed extensive labeling of villus mural cells but no tdTomato⁺ villus smooth muscle cells, indicating that the labeled PDGFRβ lineage does not directly mark mature villus smooth muscle cells (Figure S4). These validation results established that long-term lineage tracing would capture bona fide PDGFRβ-derived progeny rather than pre-existing smooth muscle cells.

**Figure 4.**
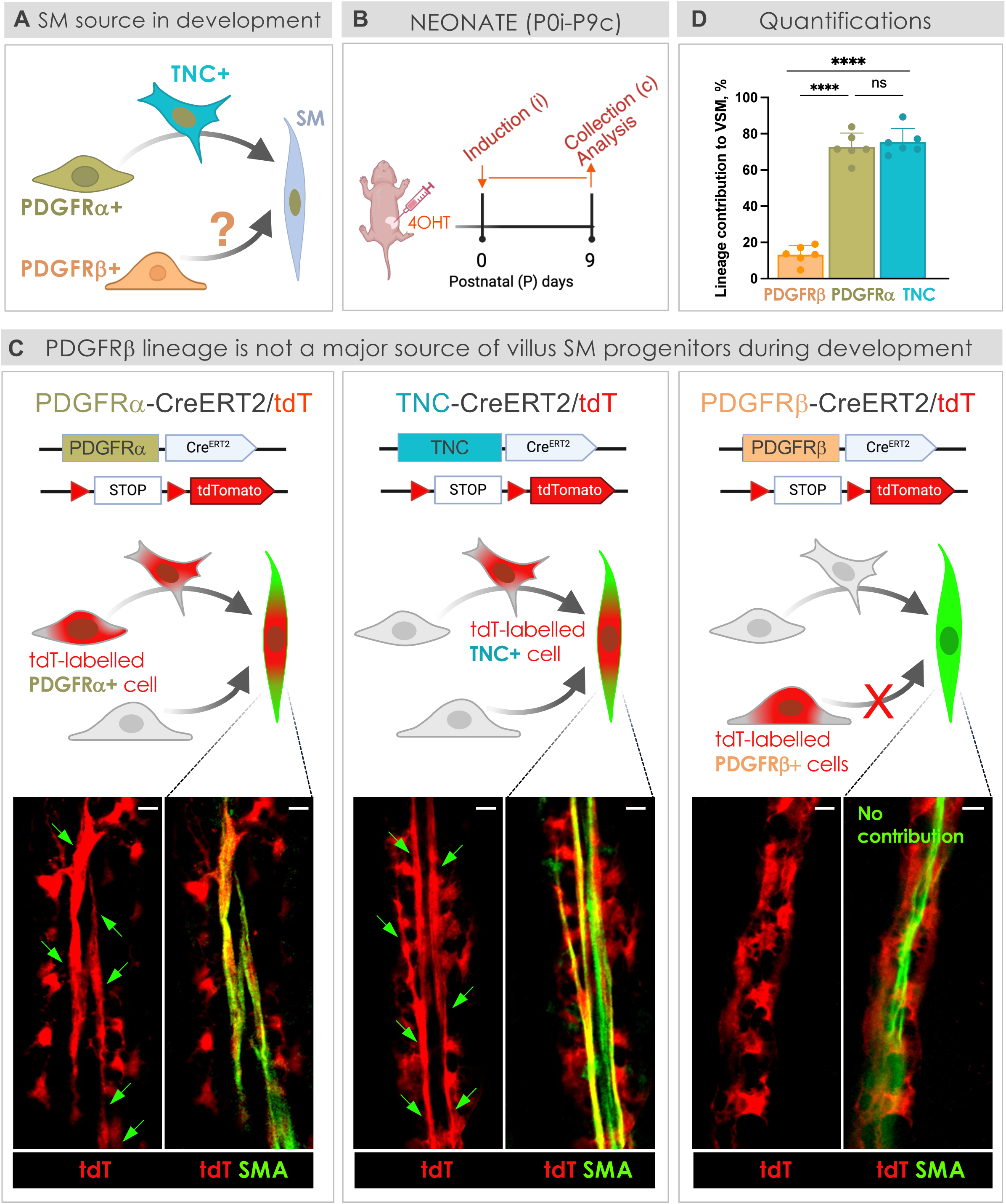
PDGFRβ lineage does not serve as a major source of villus smooth muscle progenitors during development. *See also Figure S4* (A) Cartoon depicting potential role of PDGFRβ lineage in villus SM formation. (B) Schematic for PDGFRα^+^, PDGFRβ^+^, TNC^+^ lineage tracing during the induction-collection intervals of P0i-P9c. (C) <u>Top</u>: schematic of the lineage-tracing strategy using PDGFRα-CreERT2, TNC-CreERT2, or PDGFRβ-CreERT2 crossed with Rosa26-tdTomato reporter mice. Following 4-hydroxytamoxifen-induced Cre recombination (4OHT), lineage-labeled cells permanently express tdTomato (red), enabling their contribution to villus SM to be assessed. <u>Bottom</u>: representative whole-mount confocal images showing tdTomato and αSMA (green) staining in villus SM. Arrowheads indicate tdTomato⁺/αSMA⁺ SM fibers derived from the indicated lineage. Scale bar, 10 μm. (D) Quantification of the contribution of PDGFRâ⁺, PDGFRá⁺, and TNC⁺ lineages to villus SM fibers during P0i-P9c. Contribution of <u>PDGFRβ lineage</u>: 13.23% ± 2.04%; <u>PDGFRα lineage</u>: 72.7% ± 3.13%; <u>TNC lineage</u>: 75.48% ± 3.05%. Comparisons by one-way ANOVA followed by Tukey’s multiple comparisons test. Dots indicate values from 90-120 villi/group from three independent experiments using n=6 mice/group. Data are presented as mean ± SEM. **** p <0.0001. ns, not significant.

Next, we performed lineage tracing following neonatal induction (P0–P9, Figure 4BCD). PDGFRβ-lineage cells contributed to a minor subset of villus smooth muscle fibers (13.23% ± 2.04%; Figure 4CD). In contrast, PDGFRα and TNC lineages labeled the majority of villus smooth muscle fibers during this period (72.7% ± 3.13% and 75.48% ± 3.05%, respectively; Figure 4CD). Following adult induction (P60-P69, Figure 5AB), the contribution of PDGFRβ-lineage cells was further reduced, labeling only 2.70% ± 1.04% of villus smooth muscle fibers, whereas the TNC lineage continued to contribute substantially to adult smooth muscle renewal, consistent with previous findings (20.85% ± 2.78%; Figure 5CD) (Sanketi et al. 2024).

**Figure 5.**
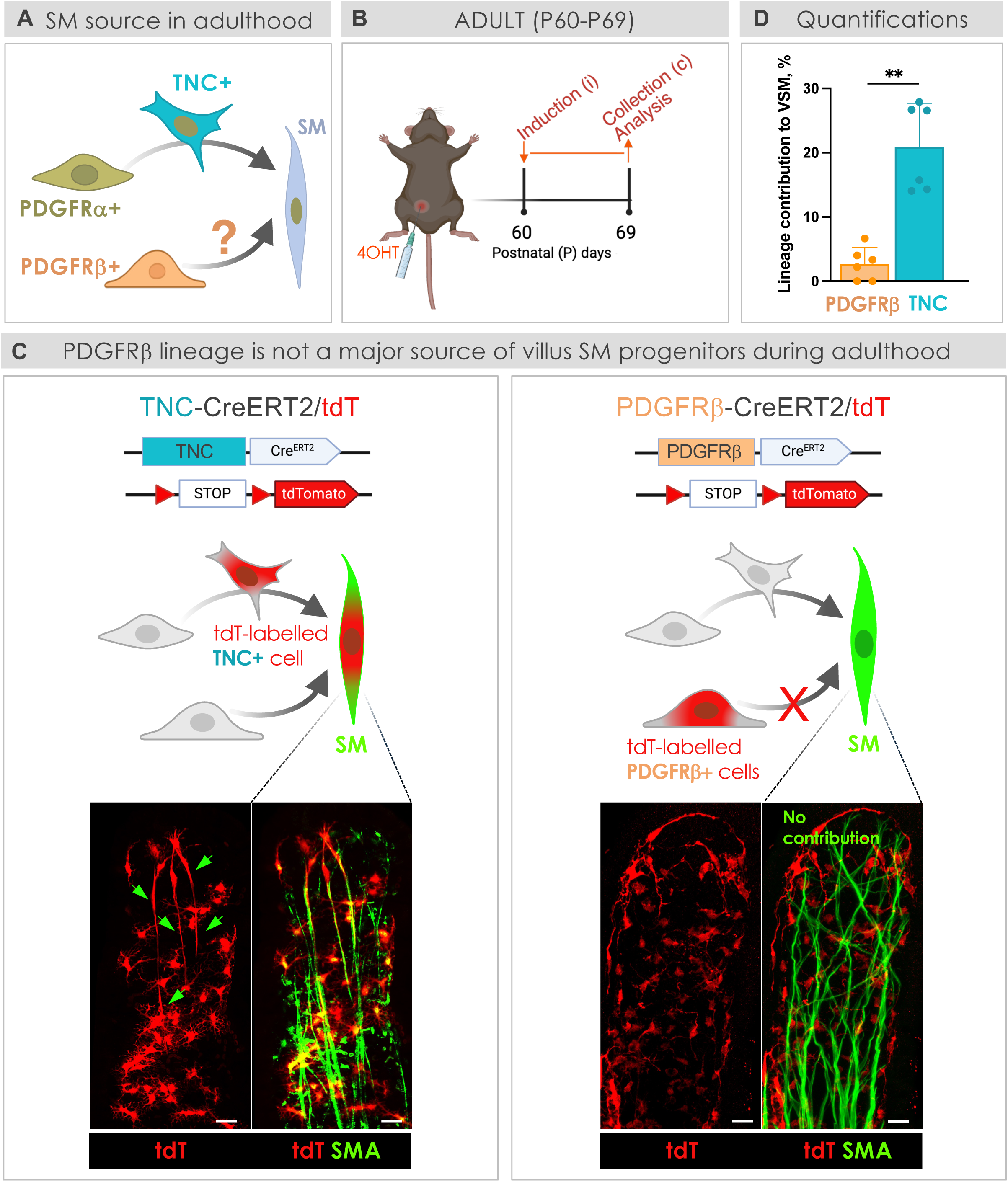
PDGFRβ lineage does not serve as a major source of villus smooth muscle progenitors in adulthood. *See also Figure S5* (A) Schematic illustrating the potential contribution of PDGFRβ⁺ lineage cells to adult villus smooth muscle renewal. (B) Experimental design for lineage tracing of PDGFRβ⁺ and TNC⁺ cells during the adult induction–collection interval (P60i–P69c). (C) <u>Top</u>: schematic of the lineage-tracing strategy. TNC-CreERT2 or PDGFRβ-CreERT2 mice were crossed with Rosa26-tdTomato reporter mice. <u>Bottom</u>: Representative whole-mount confocal images showing tdTomato (red) and αSMA (green) staining. Arrowheads indicate tdTomato⁺/αSMA⁺ villus smooth muscle fibers derived from the indicated lineage. Scale bar, 20 μm. (D) Quantification of the contribution of PDGFRβ⁺ and TNC⁺ lineages to villus SM fibers during P60i–P69c . Contribution of <u>PDGFRβ lineage</u>: 2.70% ± 1.04% and <u>TNC lineage</u>: 20.85% ± 2.78%. Statistical analysis was performed by one-way ANOVA followed by Tukey’s multiple-comparisons test. Dots indicate values from 90-120 villi/group from three independent experiments using n = 6 mice/group. Data are presented as mean ± SEM. **p = 0.0024.

Finally, we examined the fate of PDGFRβ-lineage cells in an injury-induced regeneration – a setting characterized by extensive PDGFRα^+^ mesenchymal remodeling (Jacob et al. 2022). As expected, lineage tracing following indomethacin-induced intestinal ulceration (Figure S5AB) (Jones et al.) demonstrated robust incorporation of TNC-lineage cells into regenerated villus smooth muscle (60.42% ± 5.22%)(Figure S5CE). In contrast, PDGFRβ-lineage cells showed little, if any, contribution to the regenerated smooth muscle (2.00% ± 0.98%) (Figure S5DE). Thus, PDGFRβ⁺ cells do not serve as significant smooth muscle progenitors during postnatal development, adulthood, or injury-induced repair. Instead, villus smooth muscle arises primarily from the PDGFRα lineage and regenerates through TNC⁺ intermediates (Figure S5F). These findings collectively support a non-cell-autonomous model in which PDGFRβ-lineage cells regulate villus MLC.

### PDGFRβ-lineage Notch3 promotes mesenchymal proliferation during MLC development

Expansion of the villus mesenchymal compartment is essential for generating the cellular pool required for MLC assembly. We therefore investigated whether altered mesenchymal cell growth contributed to the MLC phenotype observed following Notch3 deletion in the PDGFRβ lineage. We first examined whether increased apoptosis contributed to this defect. Western blot analysis of cleaved caspase-3 revealed no significant difference between control and Notch3ΔPdgfrβ intestines (Figure S6), indicating that impaired cell survival does not account for the loss of mesenchymal development. We therefore examined whether proliferation was affected in the absence of PDGFRβ-lineage Notch3. Strikingly, Notch3ΔPdgfrβ mice exhibited a dramatic reduction in villus stromal proliferation, as demonstrated by loss of Ki67-positive cells (Figure 6AB, and D; Control: 19.84% ± 1.14%, n = 6; Notch3ΔPdgfrβ: 1.69% ± 0.62%, n = 6). In contrast, Notch3 deletion in the PDGFRα lineage did not alter villus stromal proliferation (Figure 6 CD; Control: 11.71% ± 1.95%, n = 6; Notch3ΔPdgfrα: 8.10% ± 0.73%, n = 6). Together, these findings identify a lineage-specific requirement for Notch3 in promoting mesenchymal proliferation during MLC development.

**Figure 6.**
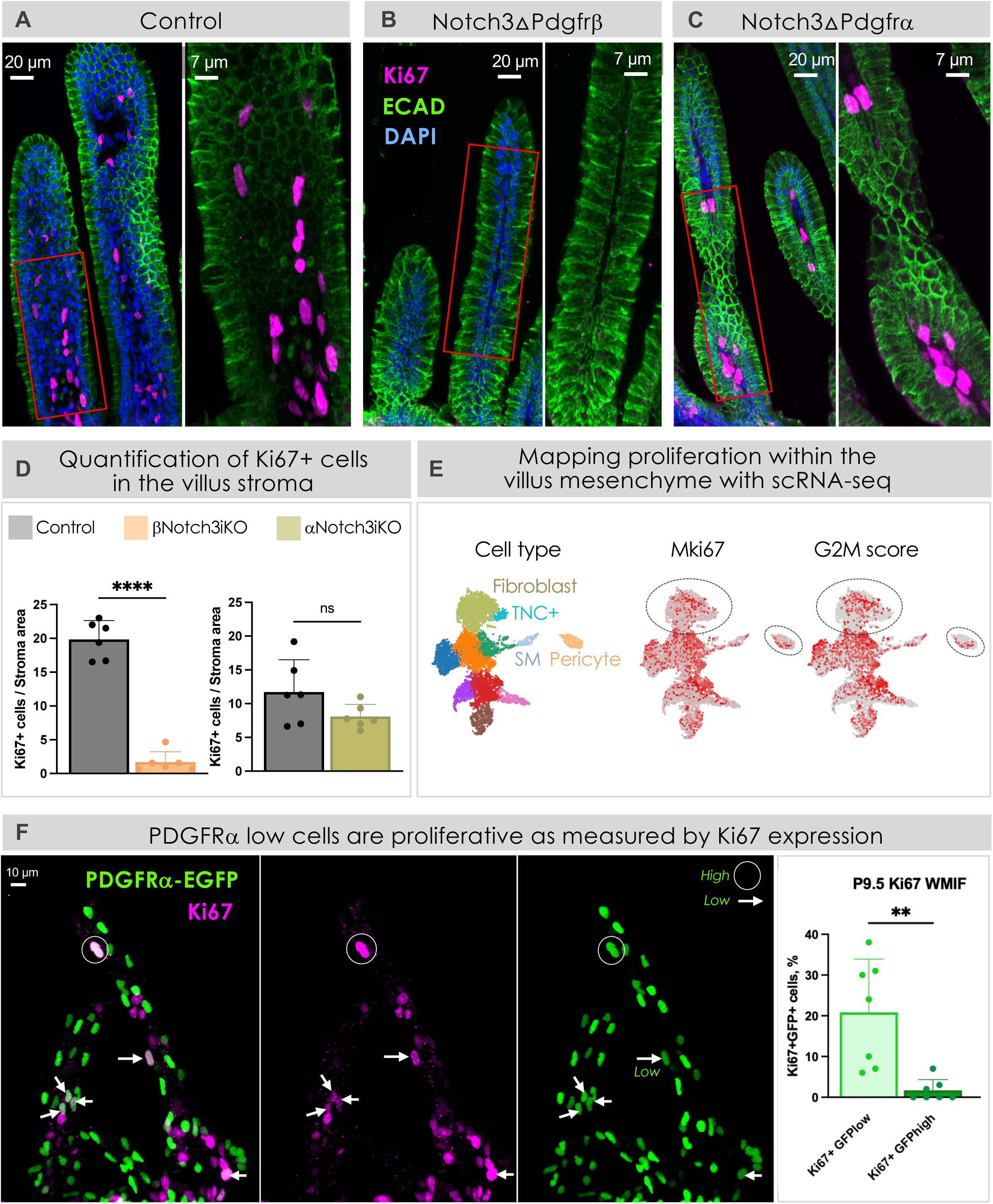
PDGFRβ-lineage Notch3 promotes mesenchymal proliferation during MLC development. *See also Figure S6* (**A-C**) Representative whole-mount confocal images showing Ki67 (magenta), E-cadherin (ECAD, green), and DAPI staining in villus smooth muscle. (**A**) Control villus, (**B**) Notch3ΔPdgfrβ, (**C**) Notch3ΔPdgfrα. Scale bar, 20 μm. Images on the right show higher-magnification views of the boxed regions (red). Scale bar, 7 μm. (D) Quantification of Ki67⁺ cells within the intestinal stroma of control, Notch3ΔPdgfrβ (βNotch3iKO), and Notch3ΔPdgfrα (αNotch3iKO) mice. Notch3 deletion in the PDGFRβ lineage, but not the PDGFRα lineage, significantly reduced stromal cell proliferation. *Control: 19.84% ± 1.14%; Notch3ΔPdgfrβ: 1.69% ± 0.62%, ****p < 0.0001. **Control: 11.71% ± 1.95%; Notch3ΔPdgfrα: 8.10% ± 0.73%, p = 0.1304 (ns). Dots indicate values from 90-120 villi/group from three independent experiments using n=6 mice/group. Data are presented as Ki67⁺ cells per stromal area (mean ± SEM). Statistical significance was determined by unpaired two-tailed Student’s *t*-test with Welch’s correction. ns, not significant. * Notch3f/f, littermate control of Notch3ΔPdgfrβ mice. ** Notch3fl/fl, littermate control of Notch3ΔPdgfrα mice. (E) UMAPs of Ki67 (Mki67) and cell cycle scores (G2M) indicating proliferative compartments in the villus mesenchyme. Fibroblast and pericyte clusters are marked by dotted circles. (F) Representative whole-mount confocal images showing Ki67+ proliferative cells with genetic reporter PDGFRα:H2b-GFP at P9. Quantification of GFP^high^Ki67⁺ (circles) and GFP^low^Ki67⁺ cells (arrows) within the intestinal stroma. GFP^low^Ki67⁺: 20.86% ± 4.93%; GFP^high^Ki67⁺: 1.71% ± 0.99%; **p = 0.0077. Dots indicate values from 16-21 villi/mice, n=2 mice. Data are presented as mean ± SEM. Statistical significance was determined by unpaired two-tailed Student’s *t*-test with Welch’s correction.

To identify the proliferating mesenchymal populations that might be affected by loss of PDGFRβ-lineage Notch3, we interrogated our developmental single-cell atlas across the P0–P9 interval. Cell cycle analysis based on Ki67 transcriptomes and G2/M-phase signatures identified several proliferative populations including PDGFRα⁺ fibroblasts and PDGFRβ^high^ pericytes (Figure 6E). Within the PDGFRα lineage, Ki67 expression was restricted to a subset of PDGFRα^low^ fibroblasts, whereas the PDGFRα^high^ fibroblasts were largely non-proliferative. The TNC⁺ myofibroblast-like population was also largely non-proliferative (Figure 6F). This pattern mirrors the fibroblast-to-myofibroblast transition, progressing from proliferation to contractile differentiation (Hinz et al. 2001; Hinz et al. 2007; Serini and Gabbiani 1999). Thus, proliferative PDGFRα⁺ smooth muscle progenitors could be depleted in PDGFRβ-lineage Notch3 mutants (P0-P9). In summary, these findings identify a PDGFRβ-lineage-specific requirement for Notch3 in maintaining the proliferative capacity of the developing villus mesenchyme, providing a mechanism by which PDGFRβ⁺ cells regulate smooth muscle development indirectly.

### PDGFRβ-lineage Notch3 promotes paracrine TGFβ signaling to PDGFRα⁺ smooth muscle progenitors

TGFβ is a well-established regulator of fibroblast proliferation, migration, and differentiation during the fibroblast-to-myofibroblast transition, setting the stage for ECM deposition (D’Urso and Kurniawan 2020). We therefore investigated the possibility that PDGFRβ⁺ cells may serve as a source of TGFβ that promotes myofibroblast differentiation of PDGFRα⁺ fibroblasts (Figure 7A). Consistent with this model, our single-cell atlas identified PDGFRβ⁺ cells as the primary source of TGFβ ligands, whereas TGFβ receptors were predominantly expressed by adjacent PDGFRα⁺ fibroblasts (Figure 7B), supporting paracrine signaling from PDGFRβ⁺ cells to smooth muscle progenitors (Figure 7A). CellChat analysis similarly predicted that the strongest TGFβ signaling interactions occurred between PDGFRβ⁺ pericytes and PDGFRα⁺ fibroblasts (Figure 7C, top). CellChat also predicted reciprocal NOTCH3 signaling between these populations, with PDGFRα⁺ fibroblasts serving as the primary signaling source to PDGFRβ⁺ cells (Figure 7C, bottom), consistent with juxtacrine activation of Notch3. Together, these findings support a model in which PDGFRβ⁺ cells receive Notch3-dependent signals from neighboring PDGFRα⁺ fibroblasts and subsequently function as a source of TGFβ, signaling back to PDGFRα⁺ smooth muscle progenitors to promote their expansion and differentiation during MLC development (Figure 7D).

**Figure 7.**
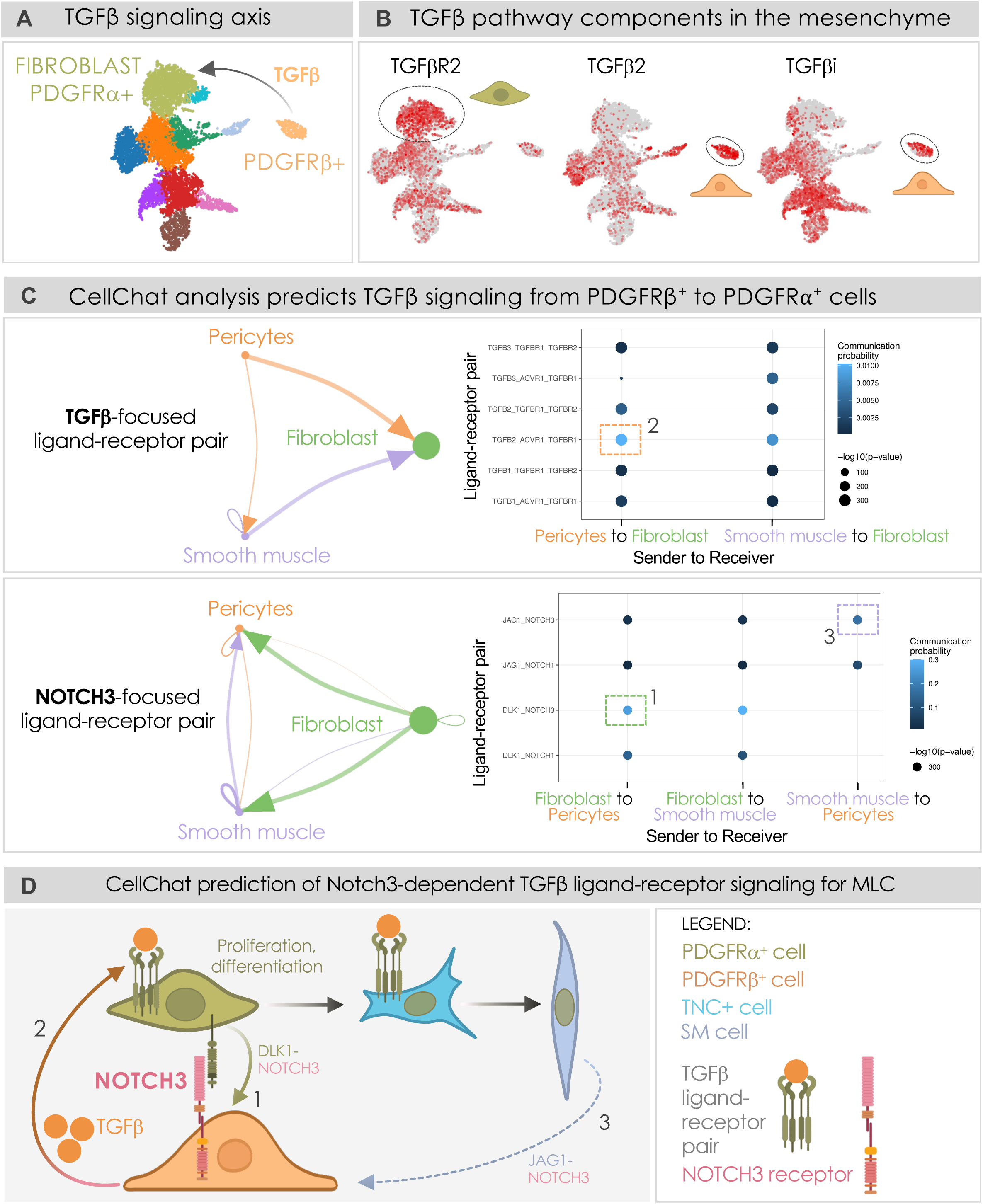
Computational analyses predict a PDGFRβ⁺→PDGFRα⁺ TGFβ signaling axis. *See also Figure S7* (A) TGFβ is a well-established regulator of fibroblast-to-myofibroblast transition, raising the possibility that PDGFRβ⁺ cells may serve as a source of TGFβ that promotes myofibroblast differentiation of PDGFRα⁺ fibroblasts. (B) UMAPs showing expression of TGFβ, TGFβ receptor 2 (TGFβR2), and TGFβ-induced) genes in P1.5 villus mesenchyme. TGFβ is enriched in the PDGFRβ⁺ pericyte cluster (circled), whereas TGFβR2 is expressed in the PDGFRα⁺ fibroblast cluster, supporting PDGFRβ⁺ cells as a potential TGFβ source for PDGFRα⁺ fibroblasts. (C) CellChat analysis identifies TGFβ and NOTCH3 signaling among major villus mesenchymal populations. CellChat analysis of TGFβ (<u>top</u>) and NOTCH (<u>bottom</u>) signaling among pericytes, fibroblasts, and villus SM. <u>Left</u>: Circle plot illustrating the inferred communication network and relative interaction strength. <u>Right</u>: Bubble plot displaying individual ligand–receptor pairs underlying the predicted interactions. Dot color represents communication probability, and dot size corresponds to statistical significance (−log10 *P* value). (D) CellChat-inspired model of reciprocal PDGFRα⁺→PDGFRβ⁺ NOTCH3 and PDGFRβ⁺→PDGFRα⁺ paracrine TGFβ signaling. In this model, PDGFRβ⁺ cells receive NOTCH3 signaling (via Dlk1-Notch3) from neighboring PDGFRα⁺ cells (1), with NOTCH3 activation predicted to regulate TGFβ production. PDGFRβ⁺-derived TGFβ then acts on PDGFRα⁺ progenitors (2, via TGFβR1) to promote their proliferation and support villus smooth muscle differentiation through a TNC⁺ intermediate “star” cell state. CellChat also predicts a villus smooth muscle→pericyte interaction through a distinct Jag1–Notch3 ligand–receptor pair, potentially providing a feedback mechanism between these populations (3).

To determine whether Notch3 regulates TGFβ signaling pathway, we performed bulk RNA-sequencing of P9 intestinal mesenchyme isolated from control and Notch3ΔPdgfrβ mice (Figure S7A). Notch3 deletion resulted in significant downregulation of TGFβ pathway genes (Figure S7B), including TGFβ-induced (TGFβi), a canonical downstream target of TGFβ (Skonier et al. 1992; Zhang et al. 2009), as well as extracellular matrix (ECM) components associated with TGFβ-mediated regulation of fibroblast-to-myofibroblast transition and ECM remodeling (Wipff et al. 2007; Bahram Yazdroudi and Malek 2023; Ansorge et al. 2017; D’Urso and Kurniawan 2020; Frangogiannis 2020). Consistent with these findings, gene set enrichment analysis (GSEA) demonstrated a significant reduction in TGFβ pathway activity in Notch3ΔPdgfrβ mesenchyme (Figure S7C). Hallmark pathways associated with cell growth and cell-cycle progression were also downregulated following loss of Notch3, consistent with our immunohistochemical analysis of Ki67 expression (Figure S7C). Together, these findings identify TGFβ as a candidate paracrine effector downstream of PDGFRβ-lineage Notch3 signaling and support a model in which PDGFRβ⁺ pericytes promote the proliferation and differentiation of neighboring PDGFRα⁺ progenitors through TGFβ signaling during MLC development.

### TGFβ supplementation rescues PDGFRβ-lineage Notch3-dependent postnatal lethality and MLC dysfunction

To determine whether activation of TGFβ signaling is sufficient to rescue the developmental defects caused by Notch3 deletion in the PDGFRβ lineage, recombinant TGFβ was administered via oral gavage to Notch3ΔPdgfrβ pups following tamoxifen induction at P0, and tissues were collected at P9 (Figure 8A). TGFβ supplementation significantly improved the growth phenotype of mutant pups, restoring body weight comparable to control levels (Figure 8B; Control: 4.76 ± 0.23 g, n = 6; Notch3ΔPdgfrβ: 3.31 ± 0.08 g, n = 6; Control + TGFβ: 4.45 ± 0.14 g, n = 6; Notch3ΔPdgfrβ + TGFβ: 4.09 ± 0.18 g, n = 6).

**Figure 8.**
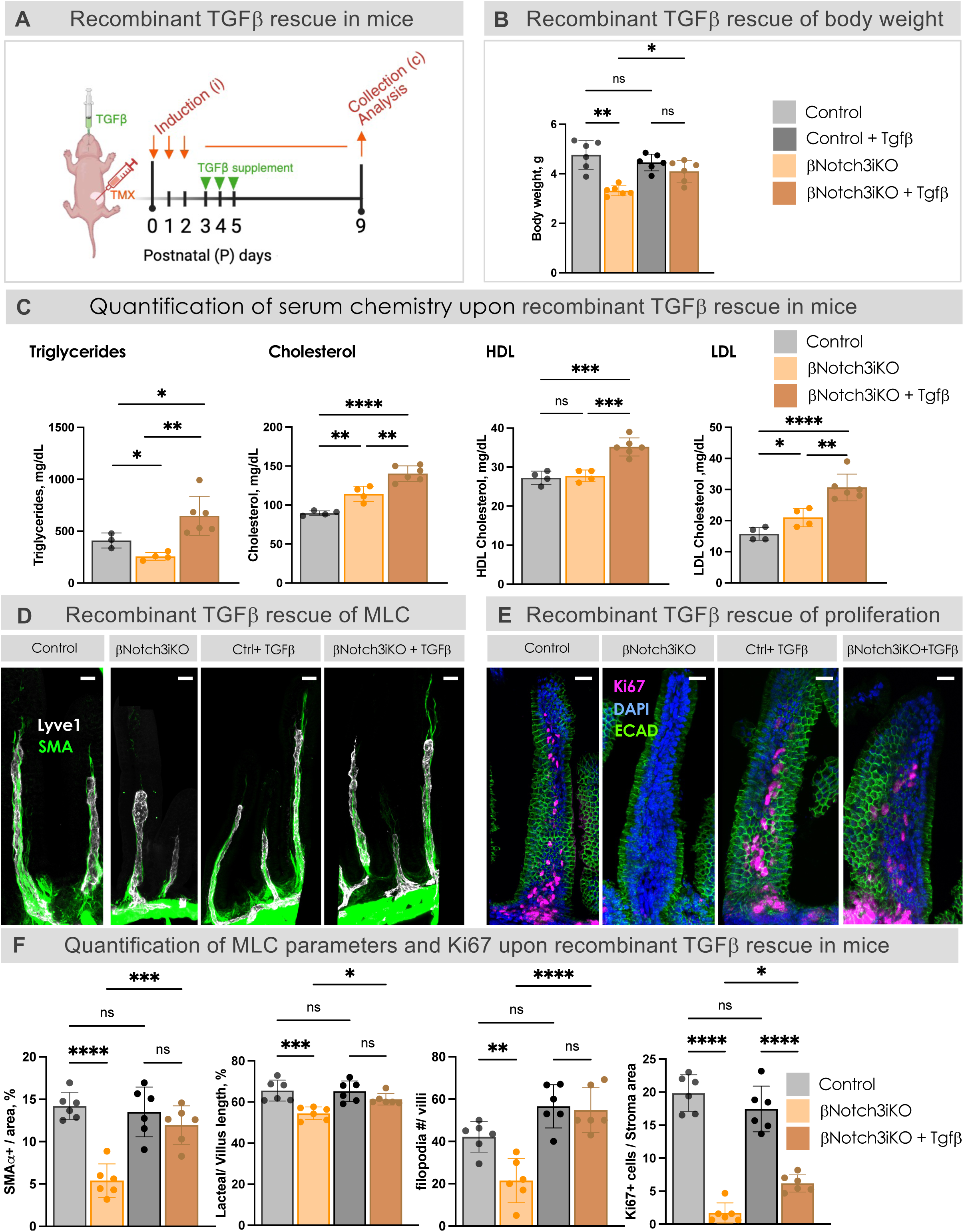
PDGFRβ-lineage Notch3 regulates MLC development through TGFβ-dependent paracrine signaling. (A) Diagram depicting the schedule of recombinant TGFβ administration to control and Notch3ΔPdgfrβ mice. (B) Body weights of control, Notch3ΔPdgfrβ (βNotch3iKO), control + TGFβ and Notch3ΔPdgfrβ(βNotch3iKO) + TGFβ. Control: 4.76 ± 0.23 g; Notch3ΔPdgfrβ: 3.31 ± 0.08 g; Control + TGFβ: 4.45 ± 0.14 g; Notch3ΔPdgfrβ + TGFβ: 4.09 ± 0.18 g. Dots indicate biological replicates, n=6, pooled from three independent experiments. Data are presented as mean ± SEM. Statistical significance was determined by one-way ANOVA followed by Tukey’s multiple-comparisons test. *p = 0.0192; **p = 0.0043; ns, not significant. (C) Serum triglycerides, cholesterol, HDL, and LDL measurements from control, Notch3ΔPdgfrβ (βNotch3iKO), and Notch3ΔPdgfrβ(βNotch3iKO) + TGFβ at P9 after oil feeding, from left to right: <u>Triglycerides</u>: Control: 409.7 ± 41.4 mg/dL, n=4; Notch3ΔPdgfrβ: 256.8 ± 19.07 mg/dL, n =4; Notch3ΔPdgfrβ + TGFβ: 647.5 ± 76.77 mg/dL, n=6; Control vs Notch3ΔPdgfrβ, *p = 0.043; Control vs Notch3ΔPdgfrβ + TGFβ, *p = 0.03; Notch3ΔPdgfrβ vs Notch3ΔPdgfrβ + TGFβ, **p = 0.0036. <u>Cholesterol:</u> Control: 89.5 ± 1.56 mg/dL, n = 4; Notch3ΔPdgfrβ: 114.3 ± 4.91 mg/dL, n = 4; Notch3ΔPdgfrβ + TGFβ: 140.3 ± 4.06 mg/dL, n = 6; Control vs Notch3ΔPdgfrβ, **p = 0.0055; Control vs Notch3ΔPdgfrβ + TGFβ, ****p < 0.0001; Notch3ΔPdgfrβ vs Notch3ΔPdgfrβ + TGFβ, **p = 0.0019. <u>HDL</u>: Control: 27.25 ± 0.85 mg/dL, n = 4; Notch3ΔPdgfrβ: 27.75 ± 0.75 mg/dL, n = 4; Notch3ΔPdgfrβ + TGFβ: 35.17 ± 0.95 mg/dL, n = 6; Control vs Notch3ΔPdgfrβ, p=0.9313(ns); Control vs Notch3ΔPdgfrβ + TGFβ, ***p = 0.0002; Notch3ΔPdgfrβ vs Notch3ΔPdgfrβ + TGFβ, ***p = 0.0003. <u>LDL</u>: Control: 15.75 ± 1.03 mg/dL, n = 4; Notch3ΔPdgfrβ: 21.0 ± 1.47 mg/dL, n = 4; Notch3ΔPdgfrβ + TGFβ: 30.67 ± 1.76 mg/dL, n = 6; Control vs Notch3ΔPdgfrβ, *p = 0.036; Control vs Notch3ΔPdgfrβ + TGFβ, ****p < 0.0001; Notch3ΔPdgfrβ vs Notch3ΔPdgfrβ + TGFβ, **p = 0.0032. Each dot represents one mouse; mice were pooled from two independent experiments, with 4 or 6 biological replicates becoming available for each group. Data are presented as mean ± SEM. Statistical significance was determined by one-way ANOVA followed by Tukey’s multiple-comparisons test. Control genotype: Notch3fl/fl. (D) Whole-mount MLCs of control, Notch3ΔPdgfrβ (βNotch3iKO), control + TGFβ and Notch3ΔPdgfrβ(βNotch3iKO) + TGFβ at P9. White is lacteal (Lyve-1); Green is aSMA (smooth muscle). Scale bar, 20 μm (E) Representative whole-mount confocal images showing Ki67(magenta), E-cadherin (green), and DAPI staining in villus smooth muscle of control, Notch3ΔPdgfrβ (βNotch3iKO), control + TGFβ and Notch3ΔPdgfrβ(βNotch3iKO) + TGFβ at P9. Scale bar, 20 μm. (F) Comparisons of the MLC parameters and Ki67 expression in control, Notch3ÄPdgfrâ (âNotch3iKO), control + TGFâ and Notch3ÄPdgfrâ(âNotch3iKO) + TGFâ at P9. <u>Villus smooth</u> <u>muscle</u>: Control: 14.21% ± 1.61%; Notch3ΔPdgfrβ: 5.40% ± 1.97%; Control + TGFβ: 13.50% ± 1.20%; Notch3ΔPdgfrβ + TGFβ: 11.96% ± 0.93%; ****p < 0.0001; ***p = 0.0003; ns. <u>Lacteal length</u>: Control: 65.55% ± 2.08%; Notch3ΔPdgfrβ: 54.46% ± 1.27%; Control + TGFβ: 65.20% ± 2.04%; Notch3ΔPdgfrβ + TGFβ: 61.31% ± 1.13%; ***p = 0.0006; *p = 0.0377; ns. <u>Percent filopodia</u>: Control: 42.19% ± 2.95%; Notch3ΔPdgfrβ: 21.50% ± 4.28%, Control + TGFβ: 56.60% ± 4.18%; Notch3ΔPdgfrβ + TGFβ: 54.82% ± 4.31%; ****p < 0.0001; **p = 0.0059; ns. <u>Ki67</u>: Control: 19.84% ± 1.14%; Notch3ΔPdgfrβ: 1.69% ± 0.62%; Control + TGFβ: 17.45% ± 1.41%; Notch3ΔPdgfrβ + TGFβ: 6.17% ± 0.53%; ****p < 0.0001; *p = 0.0184; ns. Dots indicate values from 90-120 villi/group from three independent experiments using n = 6 mice/group. Data are presented as mean ± SEM. Statistical significance was determined by one-way ANOVA followed by Tukey’s multiple-comparisons test. Control genotype: PDGFRβ-Cre/+.

To determine whether improved growth reflected recovery of intestinal absorptive function, pups were challenged with an oral lipid gavage followed by analysis of postprandial serum lipid levels. TGFβ supplementation significantly increased serum triglyceride levels in Notch3ΔPdgfrβ mice compared with untreated mutants (Figure 8C; Control: 409.7 ± 41.4 mg/dL, n = 4; Notch3ΔPdgfrβ: 256.8 ± 19.07 mg/dL, n = 4; Notch3ΔPdgfrβ + TGFβ: 647.5 ± 76.77 mg/dL, n = 6), demonstrating improved intestinal lipid uptake following TGFβ treatment. In addition, TGFβ-treated Notch3ΔPdgfrβ mice exhibited increased postprandial total cholesterol (Figure 8C; Control: 89.5 ± 1.56 mg/dL, n = 4; Notch3ΔPdgfrβ: 114.3 ± 4.91 mg/dL, n = 4; Notch3ΔPdgfrβ + TGFβ: 140.3 ± 4.06 mg/dL, n = 6), HDL (Figure 8C; Control: 27.25 ± 0.85 mg/dL, n = 4; Notch3ΔPdgfrβ: 27.75 ± 0.75 mg/dL, n = 4; Notch3ΔPdgfrβ + TGFβ: 35.17 ± 0.95 mg/dL, n = 6), and LDL levels (Figure 8C; Control: 15.75 ± 1.03 mg/dL, n = 4; Notch3ΔPdgfrβ: 21.0 ± 1.47 mg/dL, n = 4; Notch3ΔPdgfrβ + TGFβ: 30.67 ± 1.76 mg/dL, n = 6), further supporting restoration of systemic lipid transport after TGFβ supplementation.

We next examined whether TGFβ treatment corrected the structural defects in the MLC caused by loss of PDGFRβ-lineage Notch3. TGFβ supplementation improved multiple components of the complex, including lacteal length (Figure 8DF; Control: 65.55% ± 2.08%, n = 6; Notch3ΔPdgfrβ: 54.46% ± 1.27%, n = 6; Control + TGFβ: 65.20% ± 2.04%, n = 6; Notch3ΔPdgfrβ + TGFβ: 61.31% ± 1.13%, n = 6), villus smooth muscle coverage (Figure 8DF; Control: 14.21% ± 0.66%, n = 6; Notch3ΔPdgfrβ: 5.41% ± 0.81%, n = 6; Control + TGFβ: 13.50% ± 1.20%, n = 6; Notch3ΔPdgfrβ + TGFβ: 11.96% ± 0.93%, n = 6), and lacteal filopodia formation (Figure 8DF; Control: 42.19% ± 2.95%, n = 6; Notch3ΔPdgfrβ: 21.50% ± 4.28%, n = 6; Control + TGFβ: 56.60% ± 4.18%, n = 6; Notch3ΔPdgfrβ + TGFβ: 54.82% ± 4.31%, n = 6).

Consistent with a role for TGFβ in regulating mesenchymal expansion, TGFβ supplementation also partially rescued the proliferative defect observed in Notch3ΔPdgfrβ mice, increasing Ki67-positive villus stromal cells (Figure 8EF; Control: 19.84% ± 1.14%, n = 6; Notch3ΔPdgfrβ: 1.69% ± 0.62%, n = 6; Control + TGFβ: 17.45% ± 1.41%, n = 6; Notch3ΔPdgfrβ + TGFβ: 6.17% ± 0.53%, n = 6).

Together, we demonstrate that activation of TGFβ signaling is sufficient to rescue the postnatal lethality, growth impairment, and intestinal lipid absorption defects caused by loss of Notch3 in the PDGFRβ lineage. TGFβ supplementation also substantially improves MLC development and partially restores villus stromal proliferation, supporting a role for PDGFRβ-derived TGFβ signaling as an important downstream effector of Notch3-mediated mesenchymal communication. These findings establish a mechanism by which PDGFRβ⁺ cells regulate villus smooth muscle development and intestinal lipid absorption through paracrine signaling to neighboring PDGFRα⁺ mesenchymal progenitors.

### Canonical Notch signaling is dispensable for Notch3-dependent PDGFRβ-lineage regulation of MLC function

Notch3 has been reported to regulate cellular responses through both canonical RBPJ-dependent transcriptional mechanisms (Yamaguchi et al. 2008; Romay et al. 2024) and RBPJ-independent non-canonical pathways (Bhat et al. 2016). Given the established role of canonical Notch signaling in vascular smooth muscle cells (Wang et al. 2002; Nadeem et al. 2020; Baeten and Lilly 2017), we first investigated whether the defects observed following Notch3 deletion in PDGFRβ⁺ cells were mediated through disruption of canonical Notch activity by generating PDGFRβ-Cre;DNMAML^fl/fl^ (DNMAMLΔPdgfrβ) mice (Figure 9A). These mice express a dominant-negative form of Mastermind-like (DNMAML), an essential transcriptional co-activator required for canonical Notch signaling, thereby blocking RBPJ-dependent transcription downstream of all Notch receptors specifically within the PDGFRβ lineage (Shawber et al. 2019). Unexpectedly, inhibition of canonical Notch signaling in PDGFRβ⁺ cells did not reproduce the MLC defects observed following Notch3 deletion. DNMAMLΔPdgfrβ mice exhibited normal postnatal growth (Figure 9A; Control: 3.86 ± 0.10 g, n = 6; DNMAMLΔPdgfrβ: 4.03 ± 0.20 g, n = 6) and no significant abnormalities in MLC development, including normalized lacteal length (Figure 9B; Control: 60% ± 2.42%, n = 6; DNMAMLΔPdgfrβ: 56.5% ± 1.34%, n=6) or villus smooth muscle area (Figure 9B; Control: 12.64% ± 0.29%, n = 6; DNMAMLΔPdgfrβ: 11.02% ± 0.66%, n = 6). Interestingly, DNMAMLΔPdgfrβ mice displayed increased lacteal filopodia formation (Figure 8B; Control: 44.67% ± 2.85%, n = 6; DNMAMLΔPdgfrβ: 65.83% ± 7.50%, n = 6), suggesting that canonical Notch signaling may restrain aspects of lacteal growth or remodeling rather than promote neonatal MLC assembly (Suchting et al. 2007).

**Figure 9.**
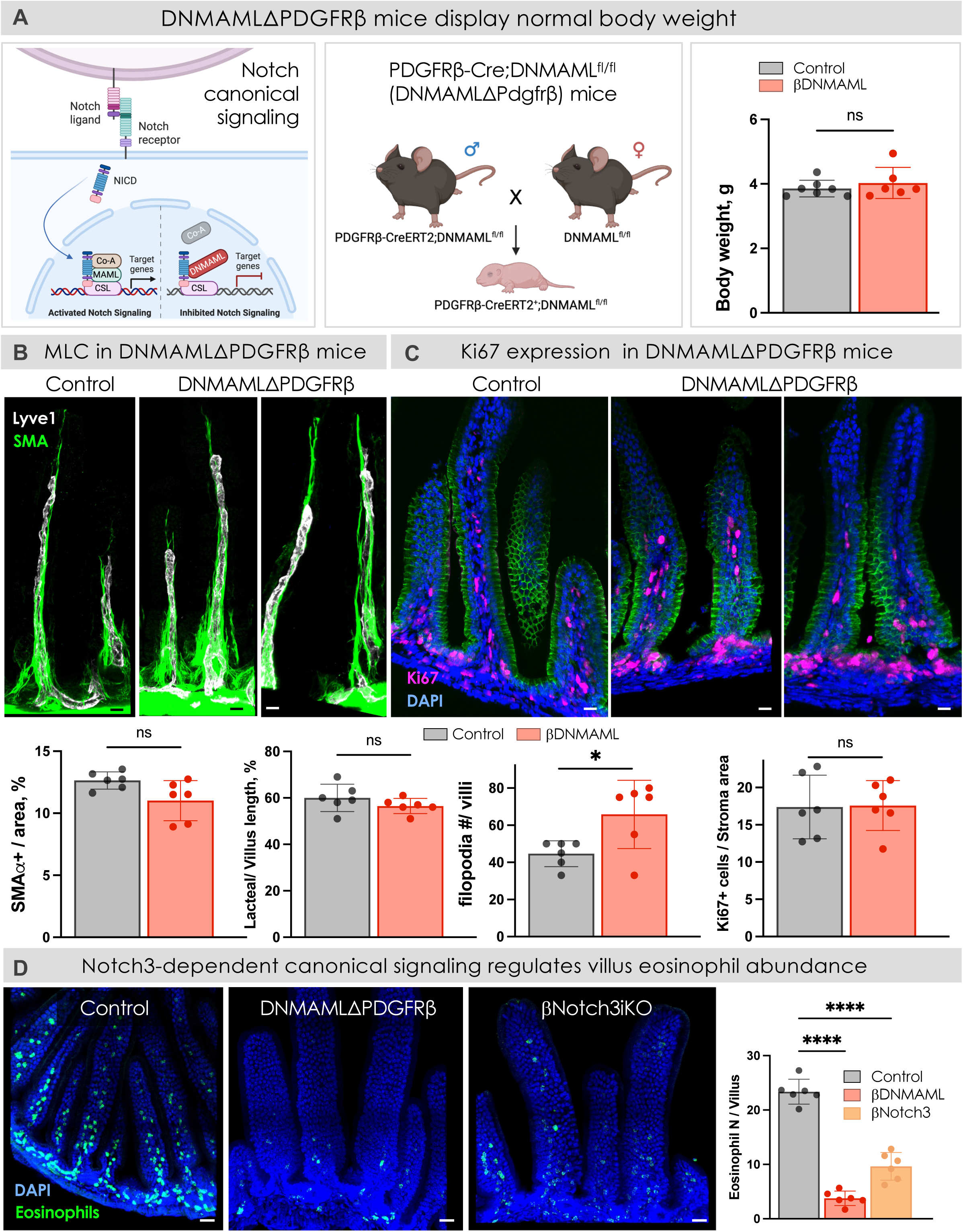
Canonical Notch is dispensable for Notch3-mediated MLC regulation. (A) <u>Left</u>: Schematic of DNMAML inhibition of canonical Notch signaling. The Notch transactivation complex consists of the Notch intracellular domain (NICD) bound to CSL on the DNA, which recruits MAML1 and coactivators (CoA) to induce transcription of Notch target genes. DNMAML-GFP fusion protein binds the NICD, blocking assembly of the activation complex and inhibiting transcription of Notch target genes. <u>Middle</u>: Schematic illustrating the breeding strategy used to generate DNMAMLΔPdgfrβ mice. <u>Right</u>: Body weight of P9 control and DNMAMLΔPdgfrβ (βDNMAML) mice. No significant difference in body weight was observed following inhibition of canonical Notch signaling in the PDGFRβ lineage: control: 3.86 ± 0.10 g; DNMAMLΔPdgfrβ: 4.03 ± 0.20 g, p = 0.4561(ns). Each dot represents one mouse, n = 6 per group, mice pooled from three independent experiments. Data are presented as mean ± SEM. Statistical significance was determined by unpaired two-tailed Student’s *t*-test with Welch’s correction. ns, not significant. Control genotype: DNMAMLfl/fl. (B) Whole-mount MLCs of control and DNMAMLÄPdgfrâ at P9 (top). White is lacteal (Lyve-1); Green is aSMA (smooth muscle). Scale bar, 20 μm. Bottom: Comparisons of the MLC parameters in control and DNMAMLΔPdgfrβ mice at P9. Villus smooth muscle area per villus: Control: 12.64% ± 0.29%; DNMAMLΔPdgfrβ: 11.02% ± 0.66%, p = 0.0596 (ns). Lacteal length normalized to villus length: control: 60% ± 2.42%; DNMAMLΔPdgfrβ: 56.5% ± 1.34%, p= 0.2423 (ns). Percent lacteals with filopodia in the jejunum: control: 44.67% ± 2.85%; DNMAMLΔPdgfrβ: 65.83% ± 7.50%, *p = 0.0363. Dots indicate values from 90-120 villi/group from three independent experiments using n=6 mice/group. Data are presented as mean ± SEM. Statistical significance was determined by unpaired two-tailed Student’s *t*-test with Welch’s correction. ns, not significant. Control genotype: DNMAMLfl/fl. (C) Representative whole-mount confocal images showing Ki67 (magenta), E-cadherin (green), and DAPI staining in villus smooth muscle. DNMAMLΔPDGFRβ mice do not recapitulate the Ki67 phenotype as in Notch3 mutants (top). Quantification of Ki67 expression per villus stroma area (bottom): control: 17.39% ± 1.74%; DNMAMLΔPdgfrβ: 17.58% ± 1.36%, p = 0.9991 (ns). Dots indicate values from 90-120 villi/group from three independent experiments using n=6 mice/group. Data are presented as mean ± SEM. Statistical significance was determined by unpaired two-tailed Student’s *t*-test with Welch’s correction. ns, not significant. Scale bar, 20 μm. Control genotype: DNMAMLfl/fl. (D) Inhibition of canonical Notch signaling in PDGFRβ causes significant reduction of intestinal eosinophils. Staining for SIGLECF (Green) and DAPI in villi of P9 Control, DNMAMLΔPDGFRβ, and Notch3ΔPDGFRβ mice. Right: quantification of the number of eosinophils per villus area: control: 23.37% ± 0.93%; DNMAMLΔPdgfrβ: 3.77% ± 0.54%; Notch3ΔPDGFRβ: 9.63% ± 1.04%, ****p < 0.0001. Dots indicate values from 90-120 villi/group from three independent experiments using n=6 mice/group. Data are presented as mean ± SEM. Statistical significance was determined by one-way ANOVA followed by Dunnett’s multiple-comparisons test. Control genotype: DNMAMLfl/fl.

We next examined whether canonical Notch signaling was responsible for the dramatic loss of villus stromal proliferation observed following Notch3 deletion. In contrast to Notch3ΔPdgfrβ mutants, DNMAMLΔPdgfrβ mice showed no reduction in Ki67-positive stromal cells (Figure 9C; Control: 17.39% ± 1.74%, n = 6; DNMAMLΔPdgfrβ: 17.58% ± 1.36%, n = 6). Thus, the proliferative defect caused by loss of Notch3 cannot be attributed to disruption of RBPJ-dependent Notch transcription, indicating that Notch3 regulates mesenchymal expansion through a distinct mechanism.

Because canonical Notch signaling has been implicated in eosinophil development (Tindemans et al. 2020), and eosinophils have recently been proposed to contribute to villus smooth muscle development through TGFβ signaling (Ignacio et al. 2022; Petrova et al. 2026), we examined whether altered eosinophil abundance could explain the Notch3ΔPdgfrβ phenotype. As expected, DNMAMLΔPdgfrβ mice exhibited a significant reduction in intestinal eosinophils (Figure 9D; Control: 23.37% ± 0.93%, n = 6; DNMAMLΔPdgfrβ: 3.77% ± 0.54%, n = 6). Notch3ΔPdgfrβ mice also showed reduced eosinophil abundance (Figure 9D; Control: 23.37% ± 0.93%, n = 6; Notch3ΔPdgfrβ: 9.63% ± 1.04%, n = 6), suggesting that eosinophil abundance is in part regulated by canonical Notch3 signaling in PDGFRβ cells. Importantly, the profound eosinophil reduction in DNMAMLΔPdgfrβ mice occurred without defects in MLC development or stromal proliferation, demonstrating that reduced eosinophil abundance alone is insufficient to account for the observed MLC phenotypes. To conclude, these experiments demonstrate that the MLC structural and functional defects caused by loss of Notch3 in PDGFRβ⁺ cells are not explained by disruption of canonical Notch signaling or eosinophils. Instead, Notch3 regulates PDGFRβ-lineage function through a distinct non-canonical mechanism and independent of eosinophils to control mesenchymal proliferation, MLC development, and lipid absorptive function.

## Discussion

Efficient intestinal lymphatic transport requires the coordinated development of lymphatic lacteals and the surrounding contractile smooth muscle that together form the MLC. While the importance of this structure for dietary lipid absorption and disease has become increasingly appreciated, the mesenchymal mechanisms that assemble and maintain this lymphatic-supporting niche have remained largely unknown. Here, we identify NOTCH3 as a central regulator of MLC development and function that coordinates communication between distinct mesenchymal lineages through genetically and mechanistically separable mechanisms (Figure 10). Whereas NOTCH3 acts cell-autonomously within the PDGFRα lineage to promote villus smooth muscle differentiation, NOTCH3 activity in PDGFRβ lineage establishes a paracrine TGFβ signaling program that supports neighboring PDGFRα^+^ smooth muscle progenitors. Together, our findings identify PDGFRβ-expressing cells as mesenchymal signaling hubs rather than smooth muscle progenitors, reveal a non-canonical mechanism of NOTCH3 function in the intestinal mesenchyme, and establish a new paradigm in which NOTCH3-directed communication between distinct mesenchymal lineages is required to build the intestinal lymphatic niche (Figure 10).

**Figure 10.**
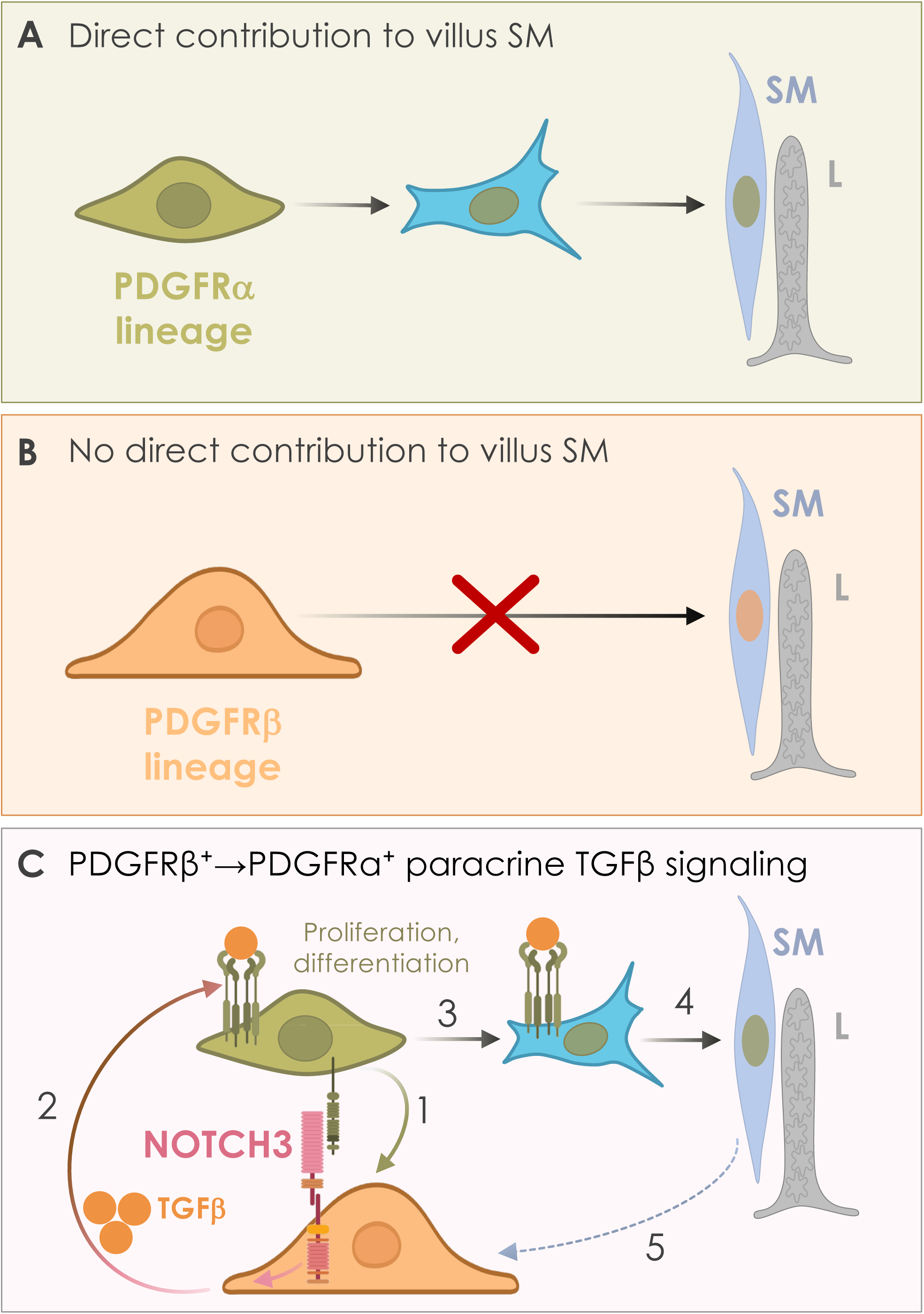
Model for PDGFR lineage-dependent Notch3 function for villus MLC. (A) PDGFRα^+^ fibroblast progenitors become mature SM, via an intermediate contractile TNC^+^ intermediate myofibroblast-like star cell (Sanketi et al. 2024; Hu et al. 2021). (B) Intestinal PDGFRβ^+^ lineage is not a major source of villus SM during development, adulthood, or injury. (C) PDGFRβ⁺ cells act as Notch3-dependent signaling hubs that drive expansion and differentiation of neighboring PDGFRα⁺ villus smooth muscle progenitors through paracrine TGFβ signaling. **1**: PDGFRβ⁺ cells receive NOTCH3 signaling from neighboring PDGFRα⁺ cells with NOTCH3 activation predicted to regulate TGFβ production. **2**: PDGFRβ⁺-derived TGFβ then acts on PDGFRα⁺ progenitors (via TGFβR1) to promote their proliferation and support villus smooth muscle differentiation through a TNC⁺ intermediate “star” cell state (**3-4**). **5**: A feedback loop may exist between villus smooth muscle cells and pericytes through a distinct Jag1–Notch3 ligand–receptor pair.

### PDGFRβ-lineage NOTCH3 indirectly supports villus smooth muscle development through paracrine TGFβ signaling

Whereas NOTCH3 acts directly within PDGFRα⁺ progenitors to promote villus smooth muscle differentiation (Sanketi et al. 2024), our findings identify a distinct indirect role for NOTCH3 in PDGFRβ⁺ cells. Rather than contributing substantially to the smooth muscle lineage, PDGFRβ⁺ cells establish a paracrine signaling environment that supports neighboring PDGFRα fibroblasts. Among the candidate pathways, TGFβ emerged as a prominent downstream effector. TGFβ is a well-established regulator of fibroblast activation (Frangogiannis 2020; Leask 2010), myofibroblast differentiation (Ansorge et al. 2017; D’Urso and Kurniawan 2020), and vascular smooth muscle development (Guo and Chen 2012) Consistent with this role, our single-cell analyses identified TGFβ2 and the canonical TGFβ target TGFβi (Lee et al. 2021; Lee et al. 2023; Skonier et al. 1992; Zhang et al. 2009) as enriched within PDGFRβ⁺ pericytes, while CellChat predicted extensive TGFβ signaling from pericytes to neighboring PDGFRα⁺ fibroblasts populations. Moreover, Notch3 deletion in the PDGFRβ lineage reduced expression of TGFβi and suppressed TGFβ pathway activity, supporting impaired TGFβ signaling as a downstream consequence of Notch3 loss.

Several observations further support this model. In contrast to Notch3 deletion in the PDGFRα lineage, loss of Notch3 in PDGFRβ⁺ cells markedly reduced stromal proliferation during early postnatal development, consistent with a diminished pool of PDGFRα⁺ smooth muscle progenitors. Importantly, restoring TGFβ signaling rescued postnatal growth, prevented lethality, and restored MLC function in Notch3ΔPdgfrβ mice. Although these findings do not exclude contributions from additional signaling pathways, they identify impaired TGFβ signaling as an important downstream consequence of Notch3 deficiency in PDGFRβ⁺ cells. Together, our findings support a model in which PDGFRβ⁺ pericytes act as signaling hubs that provide trophic support to neighboring PDGFRα⁺ progenitors through paracrine TGFβ signaling, thereby promoting formation and function of the MLC. More broadly, these results uncover an indirect mechanism by which NOTCH3 regulates smooth muscle development through intercellular communication rather than lineage contribution alone.

Our findings differ from a recent study reporting that intestinal PDGFRβ⁺ perivascular cells contribute to postnatal villus smooth muscle formation (Petrova et al. 2026). We assessed PDGFRβ⁺ lineage contribution to villus smooth muscle formation using three complementary approaches: during the neonatal period, in adulthood, and following intestinal injury induced by indomethacin treatment. We detected only a minor contribution from PDGFRβ⁺ lineage cells at P9 (13%, compared with 76% for the PDGFRα⁺/TNC⁺ lineage), and negligible contribution in adult mice or following injury (3%, compared with 60%). Thus, across the developmental and injury contexts examined, PDGFRβ⁺ lineage cells made little to no detectable contribution to villus smooth muscle compared with the substantial contribution observed from the PDGFRα⁺/TNC⁺ lineage. Several factors may account for these differing conclusions, including differences in mouse strains, genetic lineage-tracing schedules, and approaches used to quantify lineage contribution to villus smooth muscle. Importantly, both our current study and others have identified PDGFRα⁺/PDGFRβ⁺ double-positive mesenchymal cells within the intestinal stroma (Jiang et al. 2021; Han et al. 2021), potentially complicating interpretation of the contribution of distinct PDGFR+ mesenchymal populations to villus smooth muscle. Future studies using Dre/Cre intersectional genetic strategies (Han et al. 2021) to simultaneously distinguish PDGFRβ⁺ from PDGFRα⁺/PDGFRβ⁺ double-positive populations will be important to resolve the developmental potential of these heterogeneous populations with greater precision.

### Differential lineage-specific Notch3 requirement for lacteal development and function

Our findings further reveal distinct lineage-specific contributions of PDGFRα⁺ and PDGFRβ⁺ cells to lacteal development and function. Lacteals emerge around E17.5 and are functionally mature at birth to support lipid absorption from milk (Kim, Sung, and Koh 2007). Their postnatal growth and maintenance critically depend on VEGF-C signaling through VEGFR3 and its co-receptor Neuropilin-2 (NRP2), which drives lacteal sprouting, elongation, and regeneration (Tammela et al. 2008; Xu et al. 2010). Unlike lymphatic vessels in most adult tissues, whose LECs are largely quiescent, lacteals undergo continuous low-level renewal sustained by VEGF-C/VEGFR3 and DLL4–Notch signaling. VEGF-C-induced DLL4 signaling further promotes the formation of lacteal filopodia, specialized endothelial protrusions that are essential for LEC sprouting and lacteal regeneration (Bernier-Latmani et al. 2015; Xu et al. 2010). In addition, adrenomedullin–calcitonin receptor, VEGF-A/VEGFR2, and VE-cadherin signaling contribute to lacteal morphogenesis, integrity, and function (Davis et al. 2017; Zhang et al. 2018; Hägerling et al. 2018), while VEGFR3-NOTCH1 signaling is required for the postnatal formation of button junctions that enable efficient chylomicron uptake (Jannaway et al. 2023).

In the intestinal villus, VEGF-C is produced by multiple mesenchymal populations, including smooth muscle cells and PDGFRβ⁺ stroma, providing critical trophic support for neighboring LECs (Nurmi et al. 2015; Hong et al. 2020). Consistent with disruption of this supportive environment, Notch3 deletion in the PDGFRβ lineage impaired lacteal growth, resulting in significantly shorter lacteals at P9. Given that both PDGFRβ⁺ cells and smooth muscle cells contribute to villus VEGF-C expression, reduced lacteal growth may result from diminished VEGF-C production following disruption of either or both mesenchymal compartments. Interestingly, these defects are considerably more severe than those observed following Notch3 deletion in the PDGFRα lineage (Sanketi et al. 2024). In our previous study, PDGFRα-specific Notch3 deletion disrupted villus smooth muscle differentiation and MLC assembly without significantly reducing lacteal length (Sanketi et al. 2024). In contrast, PDGFRβ-specific deletion resulted in both severe villus smooth muscle and lacteal defects. Similarly, alterations in lacteal junction morphology were observed only in PDGFRβ-lineage Notch3 mutants, consistent with their impaired dietary fat absorption and associated weight loss. Together, these findings indicate that although both mesenchymal lineages cooperate to establish the MLC, they contribute through distinct Notch3-dependent mechanisms: PDGFRα-lineage cells serve as the cellular source for smooth muscle differentiation and MLC assembly, whereas PDGFRβ⁺ pericytes function as signaling coordinators that maintain the trophic environment required for MLC growth and function.

### Notch3 signaling in PDGFRβ⁺ cells is required for lipid absorption through the MLC

Dietary lipid absorption depends on the coordinated function of the MLC, in which villus smooth muscle facilitates the uptake and transport of triglyceride-rich chylomicrons into the lymphatic circulation (Sanketi et al. 2024; Hu et al. 2021; Tso et al. 2025). Consistent with the structural defects in the MLC, PDGFRβ-specific deletion of Notch3 markedly reduced circulating triglycerides, providing direct functional evidence that Notch3-dependent mesenchymal signaling is essential for intestinal lipid absorption during the neonatal period. Beyond the reduction in circulating triglycerides, total cholesterol and LDL cholesterol were increased, whereas HDL remained unchanged, indicating that Notch3 loss perturbs systemic lipid homeostasis rather than uniformly reducing circulating lipids. These changes likely reflect secondary adaptations in lipoprotein metabolism to impaired intestinal lipid uptake rather than a primary defect in cholesterol transport. Future studies will be needed to define the metabolic adaptations that accompany impaired intestinal lipid absorption. Importantly, TGFβ supplementation substantially restored circulating triglyceride levels, demonstrating that the lipid absorption defect is mechanistically linked to impaired TGFβ signaling downstream of Notch3. Given the established role of TGFβ in smooth muscle differentiation and the requirement for villus smooth muscle in lacteal-mediated lipid transport, these findings support a model in which Notch3 maintains intestinal lipid absorption by promoting TGFβ-dependent assembly and function of the MLC. Although TGFβ treatment also altered cholesterol and lipoprotein levels, these parameters reflect multiple aspects of systemic lipid metabolism and should be interpreted cautiously. Nevertheless, the robust rescue of circulating triglycerides provides compelling functional evidence that restoring the Notch3-TGFβ signaling axis is sufficient to improve intestinal lipid transport in vivo.

### Notch3 regulates villus smooth muscle primarily through non-canonical signaling

Although the canonical Notch pathway has been extensively characterized (Artavanis-Tsakonas, Rand, and Lake 1999; Kopan and Ilagan 2009), accumulating evidence indicates that Notch receptors, including Notch3, also exert important non-canonical functions. Consistent with this, CADASIL-associated NOTCH3 mutations perturb both canonical and non-canonical signaling across diverse tissues (Wang et al. 2002), highlighting the importance of non-canonical NOTCH3 activity in vivo. Canonical Notch signaling begins when a Notch ligand interacts with the extracellular domain of a Notch receptor on a juxtaposed cell (Kopan and Ilagan 2009; Siebel and Lendahl 2017; Zhou et al. 2022). Ligand-receptor interaction is followed by proteolytic cleavage events to release the Notch intracellular domain (ICD) of the receptor. Notch ICD is then trafficked to the nucleus, where it binds to the transcription factor recombination signal sequence-binding protein-JK (RBPJ/CSL) and stimulates the recruitment of Mastermind-like (MAML) and other cofactors that compose the transcriptional activation complex mediating the transcription of Notch target genes (Kopan 2012). In contrast, non-canonical Notch signaling operates independently of RBPJ and can occur through direct interactions of Notch receptors or NICDs with members of other signaling pathways, including TGFβ (Blokzijl et al. 2003; Hiratochi et al. 2007), Wnt (Hayward et al. 2005; Muñoz-Descalzo et al. 2011; Wang et al. 2007), MAPK (Carmena, Speicher, and Baylies 2006; Wang et al. 2002), and NF-κB (López-López et al. 2020).

Surprisingly, genetic inhibition of canonical Notch signaling using the DNMAML model failed to phenocopy the severe developmental defects observed in Notch3^ΔPdgfrβ^ mice. Despite marked reduction in intestinal eosinophils, DNMAMLΔPdgfrβ mice exhibited normal MLC development, stromal proliferation, and postnatal growth at P9. These findings indicate that Notch3 in PDGFRβ⁺ cells regulates MLC function largely independently of canonical Notch pathway or eosinophils.

Among candidate pathways, PDGF-B/PDGFRβ signaling has well-established roles in vascular smooth muscle and pericyte biology and functionally interacts with Notch signaling (Donovan, Abraham, and Norman 2013; Jin et al. 2008). However, we detected no changes in PDGFRβ expression in Notch3-deficient tissues at P9 (data not shown), consistent with previous findings (Baeten and Lilly 2015). Instead, our findings strongly implicate TGFβ signaling as the principal downstream effector of Notch3. TGFβ is a master regulator of vascular smooth muscle differentiation (Guo and Chen 2012; Bahram Yazdroudi and Malek 2023; Ansorge et al. 2017; D’Urso and Kurniawan 2020), and extensive evidence demonstrates that Notch and TGFβ signaling cooperate to promote the smooth muscle transcriptional program through direct interactions between NICD and SMAD2/3, resulting in activation of contractile gene expression (Tang et al. 2010). Similar cooperative signaling has been described during differentiation of multiple progenitor populations (Grieskamp et al. 2011; Kurpinski et al. 2010), while direct NICD–SMAD3 interactions have been shown to regulate transcription in other contexts (Blokzijl et al. 2003). Together with our demonstration that TGFβ supplementation rescues the developmental and functional defects of Notch3 mutants, these findings support a model in which non-canonical Notch3 signaling promotes MLC development through modulation of the TGFβ pathway rather than canonical Notch signaling itself.

### Notch3 as a central regulator of intestinal mesenchyme

Compared with the intestinal epithelium, whose regulatory networks have been extensively defined (Sato et al. 2009; McCauley et al. 2023), the genetic and molecular mechanisms governing intestinal mesenchymal growth, differentiation, and homeostasis remain poorly understood. Yet the mesenchyme is increasingly recognized as a critical regulator of intestinal development and function, providing the structural framework of the gut while establishing specialized signaling niches that support epithelial stem cells (Jacob et al. 2022; McCarthy et al. 2023), immune populations (Sun et al. 2022; Choi and Augenlicht 2024), and the lymphatic vasculature (Hu et al. 2021; Sanketi et al. 2024; Hong et al. 2020). Consequently, disruption of mesenchymal homeostasis has profound implications for intestinal disease, highlighting the need to define the pathways that govern mesenchymal specification and organization.

Our findings identify Notch3 as a central regulator of intestinal mesenchymal development. Beyond its established functions in vascular biology, we show that Notch3 orchestrates MLC formation and establishes a trophic mesenchymal niche that supports lacteal growth and function. These observations suggest that Notch3 may represent a broader organizing principle for the intestinal mesenchyme rather than a pathway dedicated solely to vascular smooth muscle differentiation.

One of the few signaling pathways previously shown to regulate intestinal mesenchymal development is the Hippo-YAP/TAZ pathway (Dang et al. 2025; Cotton et al. 2017). YAP and TAZ coordinate mesenchymal growth and patterning during gut development and are required for intestinal mesenchymal homeostasis. Intriguingly, Hippo signaling has also emerged as an important regulator of lacteal and smooth muscle development, in part through VEGF-C-dependent mechanisms (Hong et al. 2020), raising the possibility that Notch3 and Hippo signaling cooperate to coordinate expansion of the intestinal mesenchyme with assembly of the MLC. Defining the molecular interactions between these pathways will be an important direction for future studies and may reveal a broader regulatory network governing the patterning of intestinal mesenchymal development and lymphatic niche formation.

### NOTCH3-mediated mesenchymal communication in disease

Our findings have broader implications for understanding the role of NOTCH3 in vascular aging and disease. While NOTCH3 has traditionally been viewed as a cell-autonomous regulator of mural cell differentiation and contractile function, our study identifies a distinct role for NOTCH3 in coordinating communication between neighboring mesenchymal populations. This model may extend beyond the intestine. Recent single-cell transcriptomic analyses identified progressive loss of NOTCH3 expression as a molecular hallmark of vascular aging in both mice and humans, linking reduced NOTCH3 signaling to impaired vascular contractility, diminished cerebral blood flow, defective glymphatic flow, and neurodegeneration (Romay et al. 2024). Our findings raise the possibility that age-dependent decline in NOTCH3 compromises not only the intrinsic function of mural cells but also the paracrine interactions required to maintain smooth muscle homeostasis and tissue integrity. In this model, progressive loss of PDGFRβ-derived NOTCH3 signaling could impair trophic support for PDGFRα lineage-derived neighboring smooth muscle cells, thereby contributing to vascular remodeling, defective fluid transport, and tissue degeneration during aging and in NOTCH3-associated disorders such as CADASIL.

## Conclusion

In summary, our study identifies Notch3 as a central regulator of intestinal mesenchymal communication that coordinates distinct PDGFRα⁺ and PDGFRβ⁺ lineages to build the MLC. We reveal that PDGFRβ⁺ cells do not function as villus smooth muscle progenitors but instead act as Notch3-dependent signaling hubs that promote expansion and differentiation of neighboring PDGFRα⁺ smooth muscle progenitors through paracrine TGFβ signaling. These findings uncover a previously unrecognized non-canonical mechanism of Notch3 function in the heterogeneous intestinal mesenchyme and establish a new paradigm in which lineage-specific mesenchymal interactions drive lymphatic niche assembly and intestinal fat absorption. Given the emerging links between lymphatic dysfunction, vascular aging, and human disease, our findings provide a framework for understanding how disruption of mesenchymal Notch3 signaling may contribute to impaired tissue maintenance and disease progression.

## Acknowledgments

These studies would not have been possible without the contributions of past and present members of the Kurpios laboratory, with special thanks to our mouse technician, Rachel Buchanan, for her essential support. We are grateful to our collaborators for their expertise, critical discussions, and feedback on the manuscript. We thank the Cerione, Sardana, and Sethupathy laboratories at Cornell for their support and access to research resources, and are especially grateful to Drs. Marc Antonyak, Matt Zanotelli, William Lai, Shun Enomoto, Fangyu Wang, Sumanth Seetharam, and Bo Shui for their valuable contributions. Notch3- and DNMAML-flox mice were generously provided by Drs J. Kitajewski and N. Adler, and Tnc-CreERT2 mice were provided by C. M. Hao. We thank the Cornell Genomics Center for assistance with single-cell sequencing and the Cornell Bioinformatics Facility for support with computational analyses. We are grateful to the Cornell BRC Imaging Core and the Zeiss imaging team, including Dr. Jennifer Lee and Dr. Antonio Pedrosa, for microscopy support and technical expertise. We are especially grateful to Dr. Matt Thomas at the Cornell Statistical Consulting Unit (CSCU) for statistical support. We also thank the Cornell CARE program for training and assistance with our animal studies.

## Funding

National Institute of Diabetes and Digestive and Kidney Diseases R01 DK092776 (NAK); National Institute of Diabetes and Digestive and Kidney Diseases R56 DK139238 (NAK) ; National Institutes of Health 1DP2AI138242 (IDV); Cornell University Center of Vertebrate Genomics Scholarship (BDS, MM); Cornell University Center of Vertebrate Genomics Seed Grant (NAK, IDV); Cornell University College of Veterinary Medicine Graduate Research Fellowship (BDS); NIH 1S10RR025502 for Cornell Institute of Biotechnology.

## Author contributions

LH and NAK conceived and designed the study. BDS and MM performed the scRNA-seq experiments, and LH, BDS, MM, and IDV analyzed the scRNA-seq data. LH performed the experiments described in this study, including lineage tracing, immunofluorescence, genetic knockouts, bulk RNA sequencing and analysis, lipid tracing, blood lipid measurements, and indomethacin-induced injury studies. YC and TT assisted with immunofluorescence and quantitative analyses. CW contributed to tissue isolation and data interpretation. LH and NAK wrote the manuscript. All authors reviewed, provided feedback on, and approved the final manuscript.

## Data and code availability

The authors declare that all sequencing data supporting the findings of this study have been previously deposited in NCBI’s Gene Expression Omnibus with GEO series accession number <u>GSE222122</u>. All code and scripts needed to reproduce the findings of this manuscript have been previously deposited on GitHub: (https://github.com/madhavmantri/mouse_intestine_development). All other data supporting the findings in this study are included in the main article and associated files.

## Materials and correspondence

Requests for materials may be directed to (NAK)

## Conflicts

The authors declare no conflicts.

## METHODS AND KEY RESOURCE TABLE METHODS

### Animal Models

All experiments adhered to the guidelines of the Institutional Animal Care and Use Committee of Cornell University. Mice in this study were previously described: PDGFRα:H2b-GFP (JAX stock #007669) (Hamilton et al. 2003), Rosa26 CAG-tdTomato (JAX stock #007905) (Madisen et al. 2010), PDGFRβ-CreERT2 (JAX stock #030201) (Cuervo et al. 2017), PDGFRα-CreERT2(JAX stock #032770) (Chung et al. 2018), TNC-CreERT2 (a gift from C. M. Hao) (He et al. 2013), *Notch3* flox (Nadeem et al. 2020) and *DNMAML* flox (are gifts from J. Kitajewski and N. Adler) (Tu et al. 2005). All lines were maintained on a predominantly C57BL/6 genetic background. Mice were housed under pathogen-free conditions in the Cornell University animal facility on a 12-hour light/dark cycle with ad libitum access to standard rodent chow (Envigo) and water. Breeding animals were 2–6 months of age and were housed separately before mating. The day of birth was designated postnatal day 0 (P0).

Both male and female mice were included in all postnatal experiments, and sex was considered as a biological variable in all postnatal analyses. Genotyping was performed by PCR using genomic DNA isolated from neonatal toe snips collection. Mice of different genotypes and genders were randomly chosen and sorted into different procedures and measurements. When possible, littermates of different genotypes were sorted into the same procedure and measurements as pairs.

### Induction of Cre recombinase activity

Cre recombinase activity was induced for conditional gene deletion and lineage-tracing experiments by intraperitoneal administration of tamoxifen or 4-hydroxytamoxifen (4OHT; the active metabolite of tamoxifen) dissolved in peanut oil. For the 9-day neonatal conditional deletion protocol, tamoxifen was administered once daily from postnatal day 0 (P0) through P2 at a dose of 75 mg/kg per pup, and tissues were collected at P9. For the 30-day neonatal conditional deletion protocol, tamoxifen was administered once daily from P0 through P2, with additional doses administered at P9 and P18, and tissues were collected at P30. For neonatal and adult lineage-tracing experiments, 4-0HT was administered at P0 and P60, respectively; tissues were collected at P9 and P69, respectively.

For short-term PDGFRβ-lineage labeling, PDGFRβ-CreERT2; Rosa26-LSL-tdTomato mice received a single intraperitoneal injection of 4OHT (30 mg/kg) at the indicated developmental stages (P0–P2, P8–P9, or P60–P62).

### Tissue collection, immunofluorescence staining, and image acquisition

Whole-mount and cryosection immunostaining of neonatal and adult intestinal tissue was performed as previously described (Hu et al. 2021; Sanketi et al. 2024). Briefly, intestinal tissues were dissected in ice-cold 1X PBS, washed thoroughly, and fixed overnight at 4°C in either 4% paraformaldehyde (PFA) in PBS, ice-cold 100% methanol, or PAXgene® Tissue Fix. PFA-fixed tissues were washed extensively in PBS and stored at 4°C until use. Methanol-fixed tissues were stored at −20°C, whereas PAXgene-fixed tissues were processed according to the manufacturer’s instructions. For cryosectioning, tissues were cryoprotected in a graded sucrose series, embedded in OCT compound, and stored at −80°C before sectioning.

For immunofluorescence staining of frozen sections, antigen retrieval was performed using citrate-based antigen retrieval buffer, followed by permeabilization in 0.1% Triton X-100/PBS for 30 min. Sections were blocked in 3% BSA/PBS for 3 h at room temperature and incubated with primary antibodies overnight (12–16 h) at 4°C. After washing, sections were incubated with species-appropriate Alexa Fluor-conjugated secondary antibodies (Invitrogen; 1:500) together with DAPI (1:1000) for 1 h at room temperature. Slides were mounted with ProLong Gold Antifade Mountant and stored in the dark until imaging. For whole-mount immunostaining, tissues were blocked overnight (12–16 h) at 4°C in PBS containing 3% BSA, normal serum, and 0.3% Triton X-100. Samples were then incubated with primary antibodies for 24–48 h at 4°C, washed extensively in 0.3% Triton X-100/PBS, and incubated overnight with fluorophore-conjugated secondary antibodies at 4°C. Following additional washes, tissues were post-fixed overnight in 4% PFA/PBS, sectioned into 100–200 μm-thick preparations using spring scissors, and mounted in ProLong Glass Antifade Mountant. Unless otherwise indicated, primary antibodies were used at a dilution of 1:100. Anti-VE-cadherin antibody was used at 1:50. A detailed list of all antibodies used for immunofluorescence staining is provided in the Key Resources Table. Images were acquired using a Zeiss LSM 880 laser-scanning confocal microscope equipped with 40× or 63× objectives. For whole-mount preparations, adjacent fields were acquired and stitched using Zeiss ZEN software to reconstruct complete villi. Image processing, three-dimensional reconstruction, and quantitative analyses were performed using FIJI and Imaris (version 11.0).

### *In vivo* BODIPY lipid tracing assay

Dietary lipid absorption was assessed as previously described (Sanketi et al. 2024; Hu et al. 2021). Briefly, pups were fasted for 2 h before oral administration of fluorescent BODIPY™ FL C16 (Thermo Fisher Scientific, D3821). BODIPY FL C16 was dissolved in pre-warmed 20% Intralipid® emulsion (Sigma) to a final concentration of 0.4 μg/μL. Each pup received 50 μL of the BODIPY/Intralipid solution by oral gavage using a reusable 24-gauge feeding needle (Fine Science Tools, #18061-24). Following gavage, pups were returned to the dam for 2 h to maintain hydration and body temperature. Animals were euthanized by decapitation, and whole-intestine fluorescence images were acquired immediately using a Zeiss stereomicroscope.

### Serum lipid panel measurement

Serum lipid panel analysis was performed as we previously described (Sanketi et al. 2024). Briefly, after a 2-hour fast, each pup was then fed 50 ul of warmed Olive oil solution through a 24-gauge reusable feeding needle. The pups were isolated from the dam for 2 hours to avoid artifacts due to breastfeeding. A space heater and extra bedding were provided to maintain proper body temperature.

Pups were sacrificed by decapitation, and blood was collected by terminal cardiac puncture with serum separator tubes. After clotting, samples were centrifuged at 1,000 × g for 10 min to isolate serum, which was stored at −20°C until analysis. Serum triglyceride, total cholesterol, high-density lipoprotein (HDL) cholesterol, and low-density lipoprotein (LDL) cholesterol concentrations were measured by IDEXX BioAnalytics.

### Indomethacin challenge

To induce acute intestinal injury, mice received indomethacin (10 mg/kg; Sigma) by oral gavage, as previously described in established models of indomethacin-induced intestinal injury (Liang et al. 2015; Jacob et al. 2022). Control animals received an equivalent volume of vehicle (DMSO).

To examine the response of TNC⁺ and PDGFRβ⁺ mesenchymal cells following intestinal injury, TNC*-*CreERT2; Rosa26-tdTomato and PDGFRβ-CreERT2; Rosa26-tdTomato mice were first induced with 4-hydroxytamoxifen (4OHT). Two days after induction, mice were challenged with indomethacin by oral gavage and sacrificed 3 days later. Small intestines were collected for subsequent histological and immunofluorescence analyses.

### Recombinant TGFβ supplementation

Recombinant TGFβ1 (R&D, 7754-BH-025) was administered by oral gavage once daily from P4 through P6 following tamoxifen induction from P0 to P2. Recombinant TGFβ1 was diluted in 0.1% BSA/HBSS and administered at a dose of 100 μg/kg (Khounlotham et al. 2012; Wang et al. 2018). Control animals received an equivalent volume of vehicle (0.1% BSA/HBSS).

### Epithelium removal to enrich gut mesenchyme

Removal of the intestinal epithelium was adapted from previously published protocols (Sumigray, Terwilliger, and Lechler 2018; McCarthy et al. 2020). Briefly, small intestines were rapidly dissected in ice-cold PBS, washed thoroughly, and processed immediately for mesenchymal enrichment. To remove the intestinal epithelium, tissues were incubated in calcium- and magnesium-free PBS containing 10 mM EDTA at 4°C for 40 min, followed by repeated vigorous vortexing at the maximum speed until the epithelial layer was completely detached. The remaining mesenchymal tissue was gently washed in ice-cold PBS, snap-frozen in liquid nitrogen, and stored at −80°C until downstream analysis.

### Tissue processing for bulk-RNA sequencing

Total RNA was extracted from intestinal mesenchymal tissue using the RNeasy Mini Kit (QIAGEN) according to the manufacturer’s instructions, and RNA concentration and purity were assessed using a NanoDrop spectrophotometer.

### Mesenchymal RNA sequencing

Bulk RNA sequencing was performed by Plasmidsaurus (Eugene, OR) using an Illumina-based 3′-end counting RNA-seq platform. Total RNA was converted to complementary DNA (cDNA) by reverse transcription and second-strand synthesis, followed by tagmentation, library indexing, and PCR amplification. Sequencing libraries were analyzed using a bioinformatics pipeline that included FASTQ generation (BCL Convert v4.3.6; fqtk v0.3.1), quality filtering with FastP v0.24.0, alignment to the mouse reference genome using STAR v2.7, coordinate sorting with samtools v1.21, and UMI-based deduplication using UMICollapse v1.1.0. Gene expression was quantified using featureCounts (Subread v2.1.1) with strand-specific counting and summarized at the gene level. Sample normalization and principal component analysis were performed using TMM-normalized counts. Differential gene expression analysis was conducted using edgePython v0.2.5, and functional enrichment analysis was performed using GSEApy v0.12 against the MSigDB Hallmark gene sets.

### Western blotting

Small intestines were harvested, flushed with ice-cold PBS to remove luminal contents, and immediately frozen at −80°C until protein extraction. Tissues were homogenized in ice-cold T-PER™ Tissue Protein Extraction Reagent (Thermo Fisher Scientific) supplemented with protease and phosphatase inhibitor cocktails. Following centrifugation, supernatants were collected and mixed with SDS sample buffer (Bio-Rad) containing 5% β-mercaptoethanol before boiling at 95°C for 5 min.

Equal amounts of protein were separated on 4–20% Mini-PROTEAN TGX gradient gels (Bio-Rad), transferred to polyvinylidene difluoride (PVDF) membranes (Thermo Fisher Scientific), and incubated with the indicated primary antibodies. Primary antibodies included rabbit anti-cleaved Caspase-3 (Cell Signaling Technology, #9661; 1:1000) and mouse anti-β-actin (Cell Signaling Technology, #3700; 1:1000). Immunoreactive bands were detected using SuperSignal™ West Femto Maximum Sensitivity Substrate (Thermo Fisher Scientific) and imaged with a ChemiDoc XRS+ imaging system (Bio-Rad). Band intensities were quantified using FIJI.

### CellChat analysis of previously published single-cell data

Cell–cell communication analysis was performed using CellChat (v2.2.0.9001 in R) on our previously published single-cell RNA-sequencing dataset (Sanketi et al. 2024). No new single-cell sequencing data were generated for this study.

The normalized gene-expression matrix and established cell identities were used to construct a CellChat object. The mouse ligand–receptor interaction database provided with CellChat was used. Overexpressed ligands, receptors, and ligand–receptor pairs were identified using the standard CellChat workflow. Communication probabilities were inferred using default parameters.

### Image acquisition and quantification

Confocal images were taken on Zeiss LSM880 confocal microscopes. For whole-mount preparations, complete villi and lacteals were reconstructed by stitching adjacent image fields acquired with 40× or 63× objectives using Zeiss ZEN software. Image processing, background subtraction, three-dimensional reconstruction, and quantitative analyses were performed using FIJI and Imaris (version 11.0). Measurements including villus length, lacteal length, lacteal filopodia number and smooth muscle coverage were quantified using Imaris or FIJI. Lacteal junctions were quantified using Imaris software from whole-mount confocal images of intestinal lacteals. Junctions were classified into three categories based on their length, as previously described (Jannaway et al. 2023): button junctions (<4.72 μm), intermediate junctions (4.72–7.85 μm), and zipper junctions (>7.85 μm). For each lacteal, 20 endothelial junctions were analyzed, including 10 junctions from the lacteal tip to the midpoint and 10 junctions from the midpoint to the base. A total of 10–15 villi were analyzed per mouse, with six mice per experimental group. The abundance of each junctional subtype was expressed as a percentage of the total number of junctions analyzed. All z-stack datasets were analyzed in three dimensions using Imaris. For lineage-tracing experiments, individual villus smooth muscle fibers were scored for tdTomato expression based on colocalization with high αSMA immunoreactivity, as previously described (Sanketi et al. 2024).

### Statistics

Statistical analyses were performed using GraphPad Prism 10 (La Jolla, CA). Comparisons between two groups were analyzed using unpaired two-tailed Student’s *t*-tests with Welch’s correction. Comparisons among three or more groups were performed using one-way ANOVA for datasets with equal variances, followed by the Tukey post hoc test, or Welch ANOVA for datasets with unequal variances, followed by the Dunnett T3 post hoc test. Lacteal junctions were analyzed using a linear mixed-effects model (Gałecki and Burzykowski 2013; Pinheiro and Bates 2000) and t-test using Satterthwaite’s method (Zhang et al. 2018; Jannaway et al. 2023).

For imaging analyses, measurements from multiple villi were first averaged for each animal before statistical comparisons were performed, with each mouse representing a single biological replicate. Lacteal filopodia were defined as endothelial protrusions measuring ≥6 μm in length. Data are presented as mean ± SEM unless otherwise indicated. Statistical significance was defined as *P* < 0.05.

For RNA-sequencing, differential gene expression analysis was conducted using edgePython v0.2.5, and functional enrichment analysis was performed using GSEApy v0.12 against the MSigDB Hallmark gene sets.

## KEY RESOURCES TABLE

| REAGENT or RESOURCE | SOURCE | IDENTIFIER |
| --- | --- | --- |
| Antibodies |  |  |
| Mouse monoclonal anti- α-Smooth Muscle actin - FITC (SMA) | Sigma-Aldrich | Cat#F3777; RRID: AB_476977 |
| Mouse monoclonal anti- α-Smooth Muscle actin - Cy3 (SMA) | Sigma-Aldrich | Cat#C6198; RRID: AB_476856 |
| Rabbit polyclonal anti- RFP | Rockland Immunochemicals | Cat#600-401-379; RRID: AB_2209751 |
| Rabbit polyclonal anti- mouse LYVE1 | Abcam | Cat#ab14917; RRID: AB_301509 |
| Rabbit polyclonal anti- mouse Ki67 | Abcam | Cat#ab15580; RRID: AB_443209 |
| Rat monoclonal Anti-Mouse CD144 (VE-Cadherin) | BD Biosciences | Cat#555289; RRID: AB_395707 |
| Rat monoclonal Anti-Mouse Siglec-F | BD Biosciences | Cat#552125; RRID: AB_394340 |
| Rabbit polyclonal anti Cleaved Caspase-3 | Cell Signaling | Cat#9661; RRID:AB_2341188 |
| Mouse Monoclonal anti- β-Actin | Cell Signaling | Cat#3700; RRID:AB_2242334 |
| Goat anti-Rabbit IgG (H+L) Cross-Adsorbed Secondary Antibody, Alexa Fluor™ 594 | Invitrogen | Cat#A-11012; RRID:AB_141359 |
| Goat anti-Rabbit IgG (H+L) Highly Cross-Adsorbed Secondary Antibody, Alexa Fluor™ Plus 647 | Invitrogen | Cat#A32733TR; RRID:AB_2866492 |
| Goat anti-Rabbit IgG (H+L) Highly Cross-Adsorbed Secondary Antibody, Alexa Fluor™ Plus 488 | Invitrogen | Cat#A32731TR; RRID:AB_2866491 |
| Anti-rabbit IgG, HRP-linked Antibody | Cell Signaling | Cat#7074; RRID:AB_2099233 |
| Anti-mouse IgG, HRP-linked Antibody | Cell Signaling | Cat#7076; RRID:AB_330924 |
| Chemicals, peptides, and recombinant proteins |  |  |
| Tamoxifen | Sigma-Aldrich | Cat#T5648 |
| 4-hydroxy tamoxifen (4OHT) | Sigma-Aldrich | Cat#H6278 |
| Peanut oil | Sigma-Aldrich | Cat#P2144 |
| Paraformaldehyde | Sigma-Aldrich | Cat#158127 |
| O.C.T. Compound | Tissue-Tek | Cat#4583 |
| DAPI, nuclear stain | Invitrogen | Cat#D1306 |
| ProLong Gold Antifade | Thermo FisherP36930 | Cat#P36930 |
| BODIPY FL C16 (4,4-Difluoro-5,7-Dimethyl-4-Bora-3a,4a-Diaza-s-Indacene-3-Hexadecanoic Acid) | Thermo Fisher | Cat#D3821 |
| Intralipid, 20% | Sigma-Aldrich | Cat#I141 |
| Olive oil | Sigma-Aldrich | Cat#O1514 |
| T-PER™ Tissue Protein Extraction Reagent | Thermo Scientific | Cat#78510 |
| Precision plus protein Dual color standards | Biorad | Cat#1610374 |
| 4–20% Mini-PROTEAN® TGX™ Precast Protein Gels, 10-well, 50 µl | Biorad | Cat#4561094 |
| 10x Tris/Glycine/SDS buffer | Biorad | Cat#1610732 |
| 10x Tris/Glycine Buffer | Biorad | Cat#1610734 |
| 2x Laemmli Sample Buffer | Bio-Rad | Cat#1610737 |
| 2-Mercaptoethanol | Sigma-Aldrich | Cat#M3148 |
| 1X PBS, w/o calcium and magnesium, PH 7.4 ± 0.1 | Corning | Cat#21-040-CV |
| EDTA (0.5 M), pH 8.0 RNase-free | Life Technologies | Cat#AM9260G |
| Bovine serum albumin | Sigma-Aldrich | Cat#A3059 |
| Goat serum | Thermo Scientific | Cat#16210072 |
| Triton™ X-100 | Sigma-Aldrich | Cat#TX1568-1 |
| Tween-20 | FisherBiotech | Cat#BP337500 |
| FocusClear™ | Celexplorer | Cat#FC-101 |
| Halt™ Protease and Phosphatase Inhibitor Cocktail | Thermo Scientific | Cat#78447 |
| Dimethyl Sulfoxide (DMSO) | Sigma-Aldrich | Cat#D2650 |
| SuperSignal™ West Femto Maximum Sensitivity Substrate | Life Technologies | Cat#34095 |
| PAXgene Tissue FIX and Stabilizer | QIAGEN | Cat#765312, 765512 |
| Critical commercial assays |  |  |
| Serum lipid measurement | IDEXX Bioanalytics | Rodent Lipid Panel (1-6) |
| Experimental models: Organisms/strains |  |  |
| PDGFRα:H2b-GFP | Jackson Labs (Hamilton et al. 2003) | Strain #007669 |
| Rosa26 CAG-tdTomato | Jackson Labs | Strain #007905 |
|  | (Madisen et al. 2010) |  |
| PDGFRβ-CreERT2 | Jackson Labs<br>(Cuervo et al. 2017) | Strain #030201 |
| PDGFRα-CreERT2 | Jackson Labs<br>(Chung et al. 2018) | Strain #032770 |
| TNC-CreERT2 | Gift from Dr. Chuan-Ming Hao, Fudan University / Vanderbilt University<br>(He et al. 2013) | N/A |
| Notch3 flox | Gift from Drs. Jan Kitajewski and Naiche Adler, University of Illinois<br>(Nadeem et al. 2020) | N/A |
| DNMAML flox | Gift from Drs. Jan Kitajewski and Naiche Adler, University of Illinois<br>(Tu et al. 2005) | Strain #032613 |

| Software and algorithms |  |  |
| --- | --- | --- |
| Zen microscopy suite | Zeiss | RRID:SCR_013672 |
| Prism Graphpad 10 | Graphpad | RRID: SCR_005375 |
| ImageJ /Fiji | NIH | RRID: SCR_002285 |
| Imaris 11 | Bitplane | RRID: SCR_007370 |
| CellChat (v2.2.0.9001) | Jin et al.(Jin, Plikus, and Nie 2025) | https://github.com/jinworks/CellChat |

| Other |  |  |
| --- | --- | --- |
| Zeiss Confocal Microscope, 880 | Zeiss | RRID: SCR_020925 |
| ChemiDoc MP Imaging System | Biorad | RRID:SCR_019037 |

## Notes

https://github.com/madhavmantri/mouse_intestine_development

### Competing Interest Statement

The authors have declared no competing interest.

