## Supplemental Figures and Figure Legends 1-7 for "A NON-CANONICAL ROLE FOR NOTCH3 IN BUILDING THE INTESTINAL LYMPHATIC NICHE"

**Figure S1. Expression of genes defining the villus pericyte cluster at P1.5**

*Relates to Figure 1 in Main Text*

UMAP plots showing the expression of canonical and novel markers associated with intestinal pericytes at P1.5.

*Relates to Figure 1 in Main Text*

UMAP plots of villus pericyte cluster defining genes from single cell RNA-sequencing (P1.5)

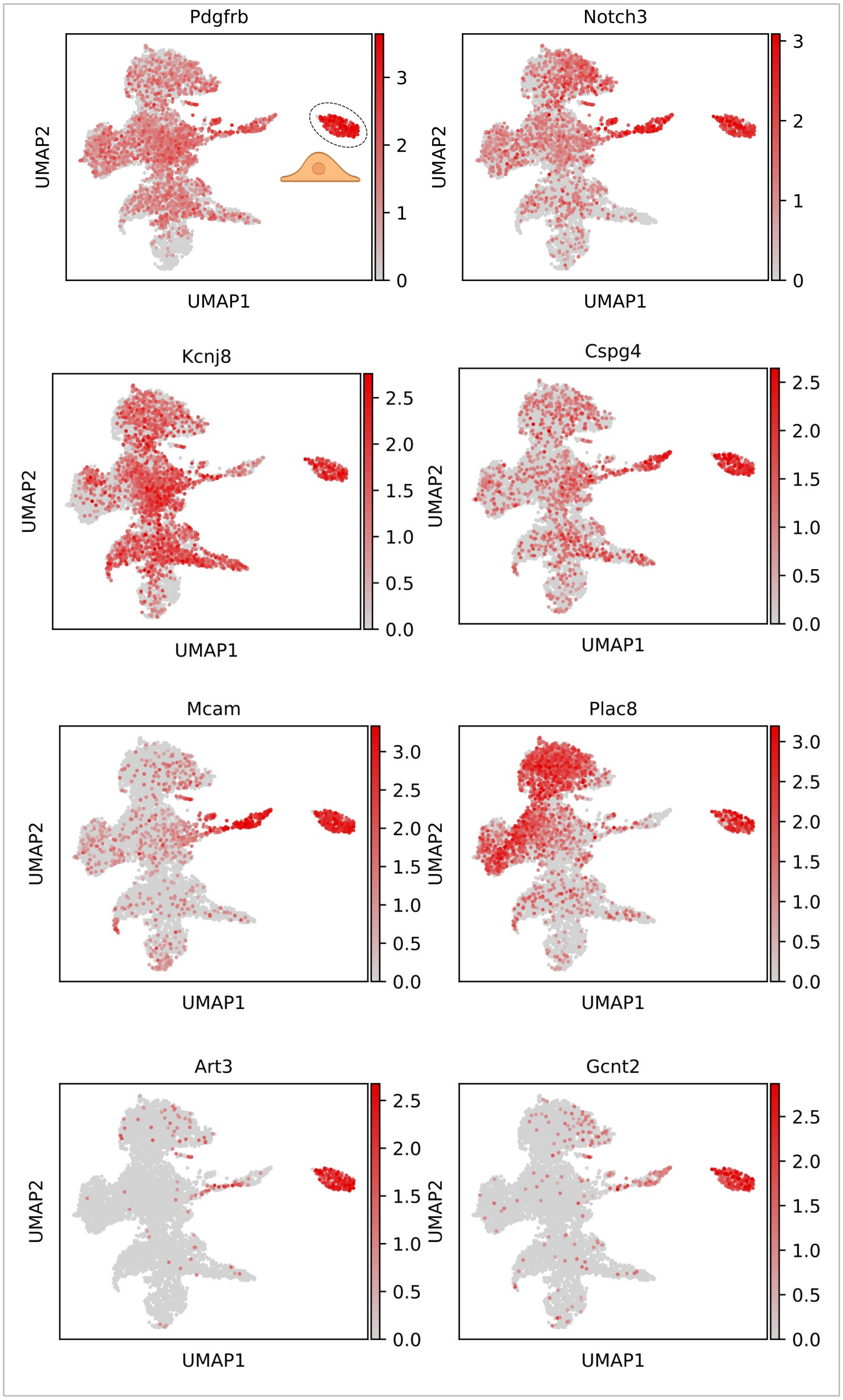

**Figure S2. PDGFRβ-lineage Notch3 loss disrupts postnatal growth and survival**

*Relates to Figure 2 in Main Text*

(**A**) A pair of mice showing significant growth retardation of Notch3ΔPdgfrβ versus control at P30.

(**B**) Left: Diagram depicting the generation of Notch3ΔPdgfrβ mouse and PDGFRβ+ cell-specific depletion of Notch3 in newborn pups and analyses at P30. Right: Mortality rates of control and Notch3ΔPdgfrβ mice at P30. P30 control: 0.00 % ± 0.00 %; P30 Notch3ΔPdgfrβ: 58.25% ± 14.48 %, * p=0.0276. Each dot indicates a group of mice, n = 3 (control) or n = 4 (Notch3ΔPdgfrβ) mice pooled from three and four independent experiments, respectively. Data are presented as mean ± SEM. Statistical significance was determined by unpaired two-tailed Student's t-tests with Welch's correction. Control genotype: PDGFRβ-Cre/+; Rosa26-tdTomato

*HUANG et al. SUPP FIGURE 2*

*Relates to Figure 2 in Main Text*

**A** Notch3 is required in PDGFRβ$^{+}$ cells for postnatal growth

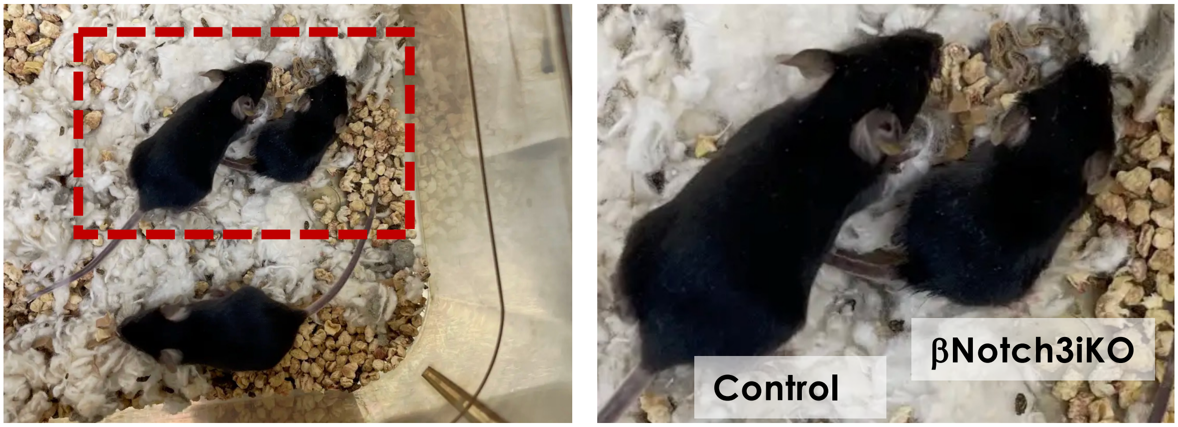


**B** Loss of Notch3 in PDGFRβ-lineage results in postnatal lethality

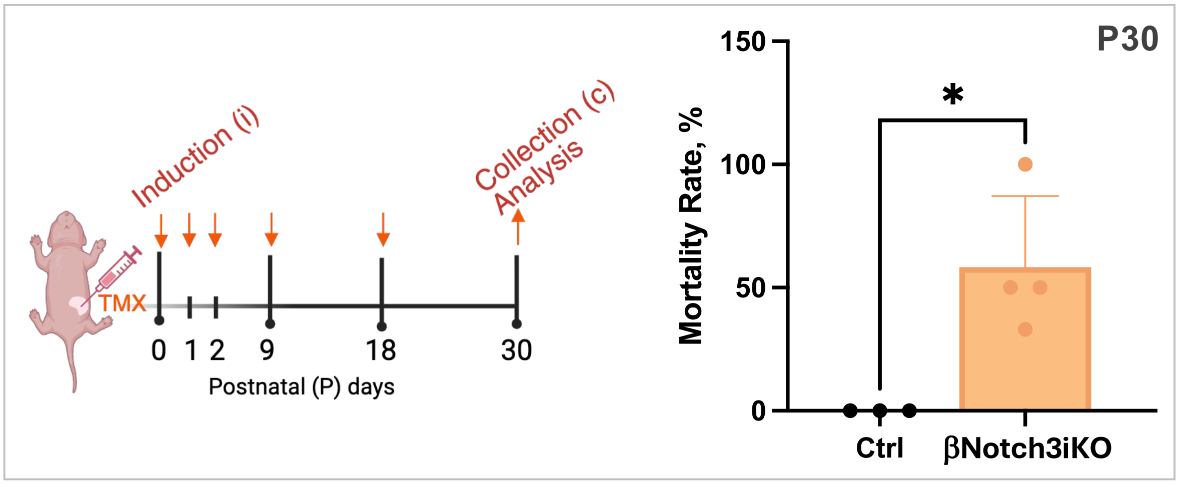

**Figure S3. Impaired lipid uptake upon loss of Notch3 in PDGFRβ lineage**

*Relates to Figure 3 in Main Text*

(**A**) Schematic for feeding olive oil and downstream lipid panel analysis in mouse pups using IDEXX lab testing.

(**B**) Measurements of triglycerides, cholesterol, HDL, and LDL from control and Notch3ΔPdgfrβ pups after oil feeding. Triglycerides: Control: 409.7 ± 41.4 mg/dL; Notch3ΔPdgfrβ: 256.8 ± 19.07 mg/dL, *p = 0.0473. Total cholesterol: Control: 89.5 ± 1.56 mg/dL; Notch3ΔPdgfrβ: 114.3 ± 4.91, *p = 0.0112. HDL: Control: 27.25 ± 0.85 mg/dL; Notch3ΔPdgfrβ: 27.75 ± 0.75 mg/dL, p = 0.6756 (ns). LDL: Control: 15.75 ± 1.03 mg/dL; Notch3ΔPdgfrβ: 21.0 ± 1.47 mg/dL, *p = 0.03. Each dot represents one mouse; mice were pooled from two independent experiments, with 4 biological replicates becoming available for each group. Data are presented as mean ± SEM. Statistical significance was determined by unpaired two-tailed Student's t-tests with Welch's correction. Control genotype: Notch3fl/fl.

*HUANG et al. SUPP FIGURE 3*

*Relates to Figure 3 in Main Text*

**A** Serum chemistry analysis in Notch3ΔPdgfrβ

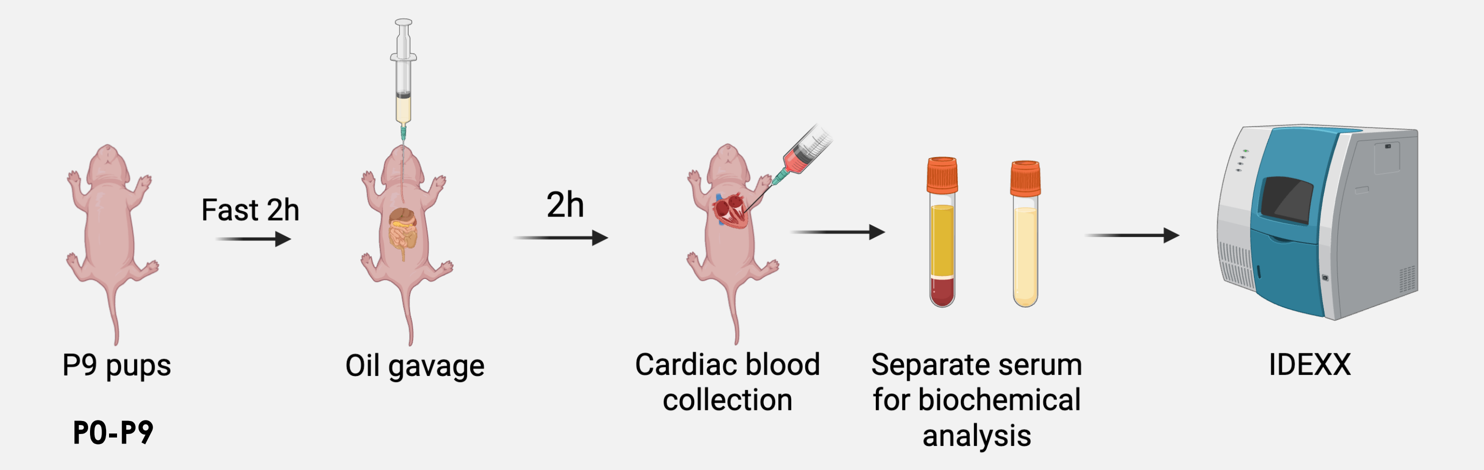


**B** Quantification of serum chemistry

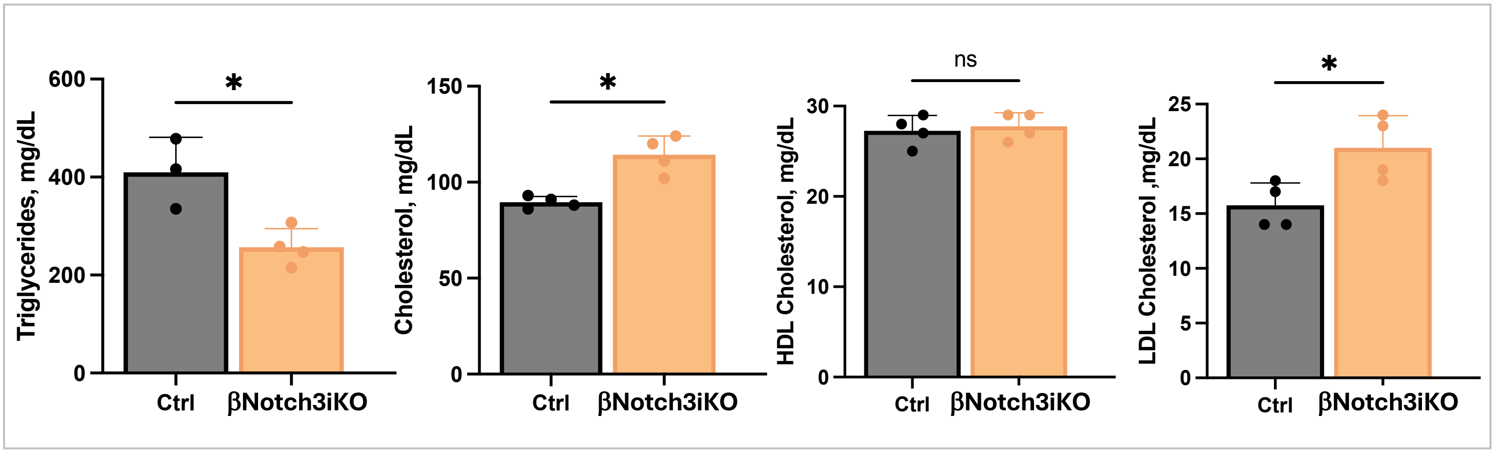

**Figure S4 Intestinal villus smooth muscle cells do not express PDGFRβ**

*Relates to Figure 4 in Main Text*

Control lineage tracing using PDGFRβ-CreERT2;Rosa26-tdTomato mice following short induction-to-collection intervals: (**A**) P0i-P3c, (**B**) P8i-P9c, and (**C**) P60i-P62. Whole-mount immunostaining of αSMA (green) and tdTomato (red). No $\alpha SMA^+$/$tdTomato^+$ double-positive villus SM cells were detected at any time point, demonstrating that differentiated villus SM cells do not express PDGFRβ. Scale bar, 20 µm. Boxed regions are shown at higher magnification on the right. Scale bar, 5 µm.

*HUANG et al. SUPP FIGURE 4*

*Relates to Figure 4 in Main Text*

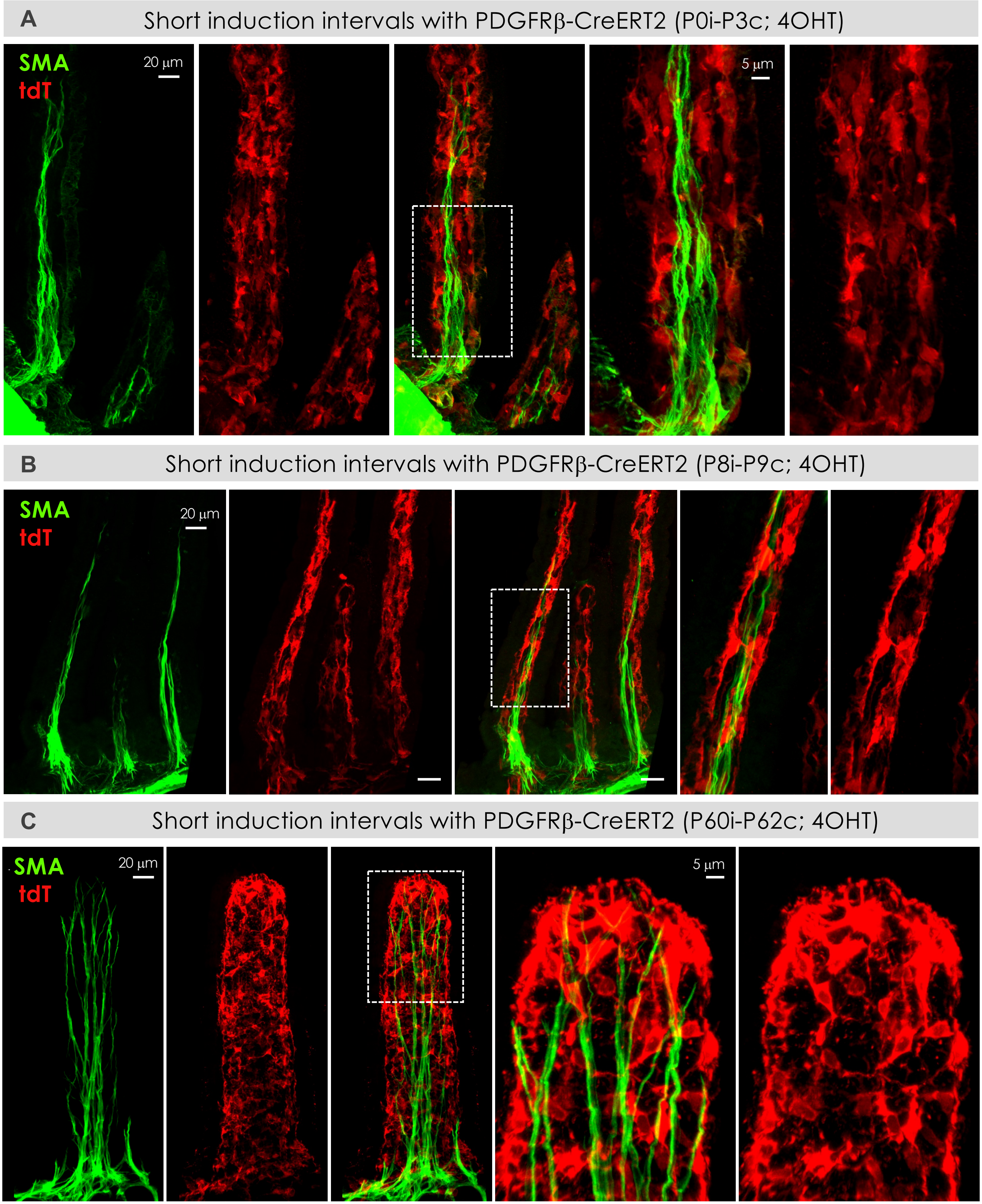

**Figure S5. PDGFRβ lineage does not serve as a major source of villus smooth muscle progenitors upon injury**

*Relates to Figure 5 in Main Text*

(**A**) Schematic for indomethacin treatment in adult mice.

(**B**) Histology of vehicle- (top) and indomethacin (bottom)-treated jejunum. Scale bar, 100 µm

(**C, D**) Representative whole-mount confocal images of TNC-CreERT2; Rosa26-tdTomato (**C**) and PDGFRβ-CreERT2; Rosa26-tdTomato (**D**) intestines showing tdTomato (red) and αSMA (green) staining. Arrows indicate $tdTomato^{+}/\alpha SMA^{+}$ villus smooth muscle fibers derived from the indicated lineage. Scale bar, 15µm.

(**E**) $PDGFR\beta^{+}$ and $TNC^{+}$ lineage tracing contributions to SM fibers in response to indomethacin challenge. Contribution of <u>PDGFRβ lineage</u>: 2.00% ± 0.98% and <u>TNC lineage</u>: 60.42% ± 5.22%. Dots indicate values from 90-120 villi/group from three independent experiments using n=6 mice/group. Data are presented as mean ± SEM. Statistical significance was determined by unpaired two-tailed Student's t-tests with Welch's correction. **** $p < 0.0001$.

(**F**) Proposed model of villus SM regeneration in which $TNC^{+}$ progenitors, but not PDGFRβ-lineage cells, are the predominant source of new SM after injury.

*HUANG et al. SUPP FIGURE 5*

*Relates to Figure 5 in Main Text*

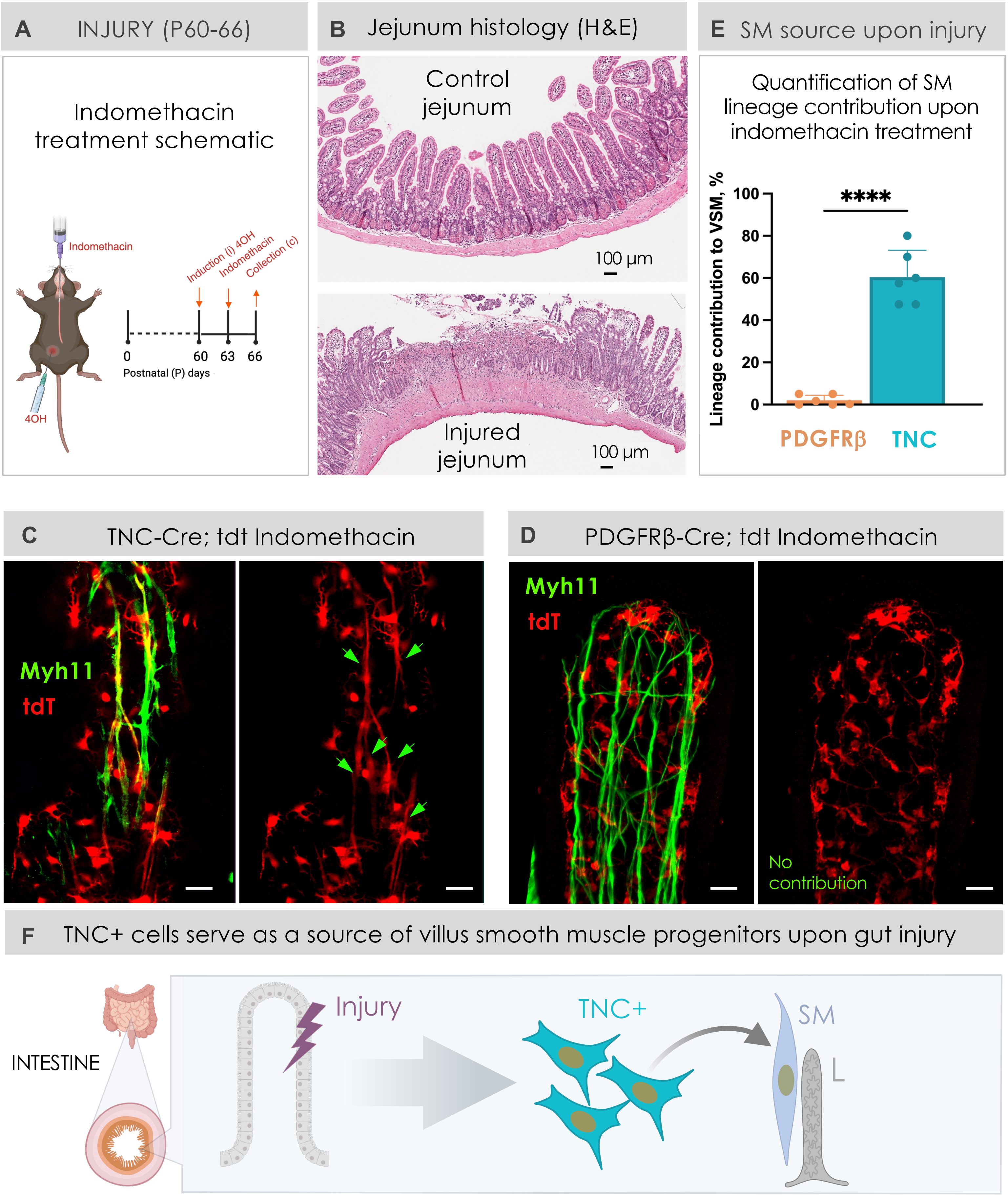

**Figure S6. Apoptosis is not affected by the loss of Notch3 from PDGFRβ lineage at P0-P9**

*Relates to Figure 6 in Main Text*

(**A**) Western blot showing cleaved Caspase-3 and β-actin expression in small intestinal tissue from P9 Control (gray), Notch3ΔPdgfrβ (orange), and Notch3DNMAML (red) mice. β-Actin served as a loading control. Control genotype: PDGFRβ-Cre/+.

(**B**) Quantification of cleaved Caspase-3 normalized to β-actin. No significant differences in cleaved Caspase-3 protein levels were detected among the three groups. n=3 mice per group. Data are presented as mean ± SEM. Statistical analysis was performed by one-way ANOVA followed by Tukey's multiple-comparisons test. ns, not significant.

*HUANG et al. SUPP FIGURE 6*

*Relates to Figure 6 in Main Text*

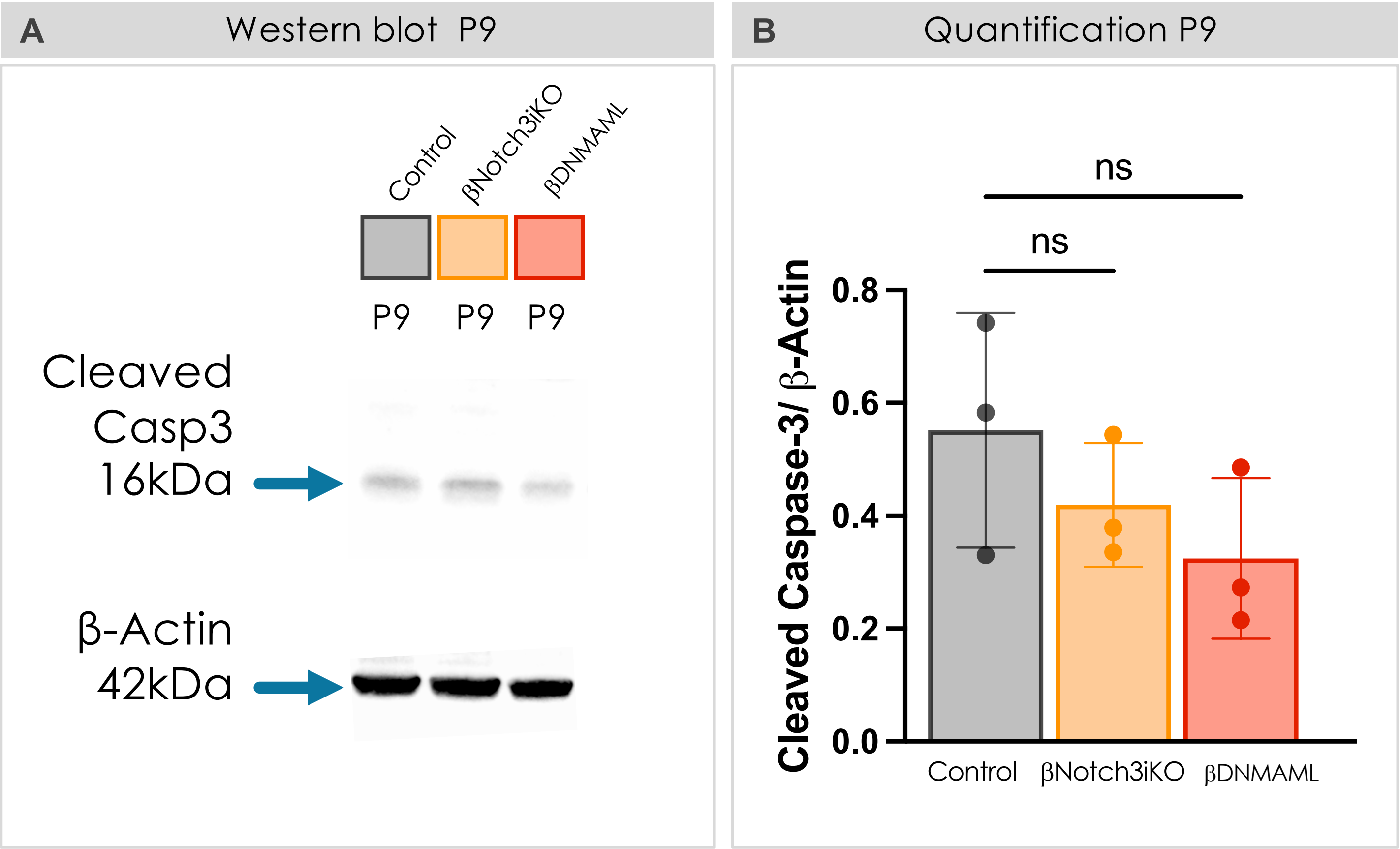

**Figure S7 Bulk RNA-sequencing of the gut mesenchymal compartment**

*Relates to Figure 7 in Main Text*

(**A**) Diagram depicting the procedure of intestinal epithelium removal and mesenchymal enrichment for bulk RNA-sequencing of the gut mesenchyme.

(**B**) Heatmap of selected TGFβ signaling, extracellular matrix (ECM), and Notch3 genes in bulk RNA-seq of P9 intestinal mesenchyme from control and Notch3ΔPdgfrβ (βNotch3iKO) mice (n = 3 per group). Expression values are displayed as row-scaled z-scores.

(**C**) Gene set enrichment analysis (GSEA) of bulk RNA-seq data from intestinal mesenchyme of control and Notch3ΔPdgfrβ mice at P4 and P9. Bar plots show the normalized enrichment scores (NES) of significantly downregulated MSigDB Hallmark gene sets in Notch3ΔPdgfrβ mice relative to controls. Red boxes: TGFβ pathway hallmark genes; Green boxes (cell growth and cell cycle proliferation hallmark genes). Control genotype: PDGFRβ-Cre/+.

*HUANG et al. SUPP FIGURE 7*

*Relates to Figure 7 in Main Text*

**A** Intestinal mesenchyme-targeted bulk RNA-sequencing

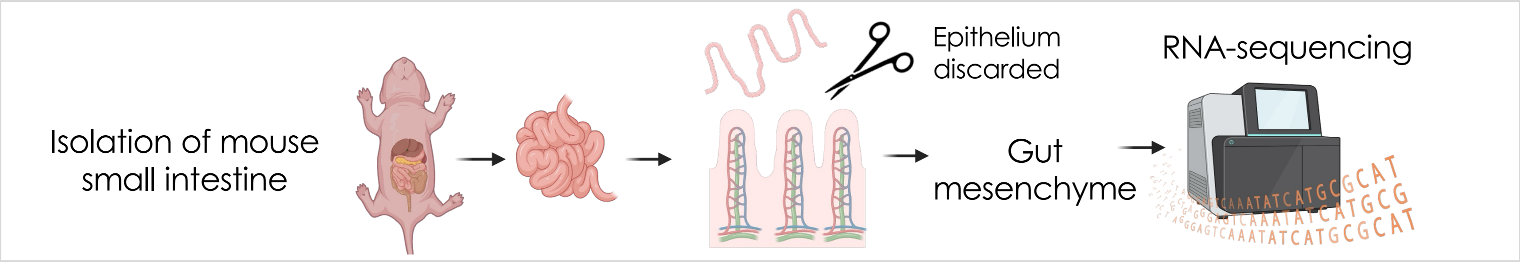


**B** Mesenchyme RNA-seq reveals Notch3-dependent TGFβ and ECM programs

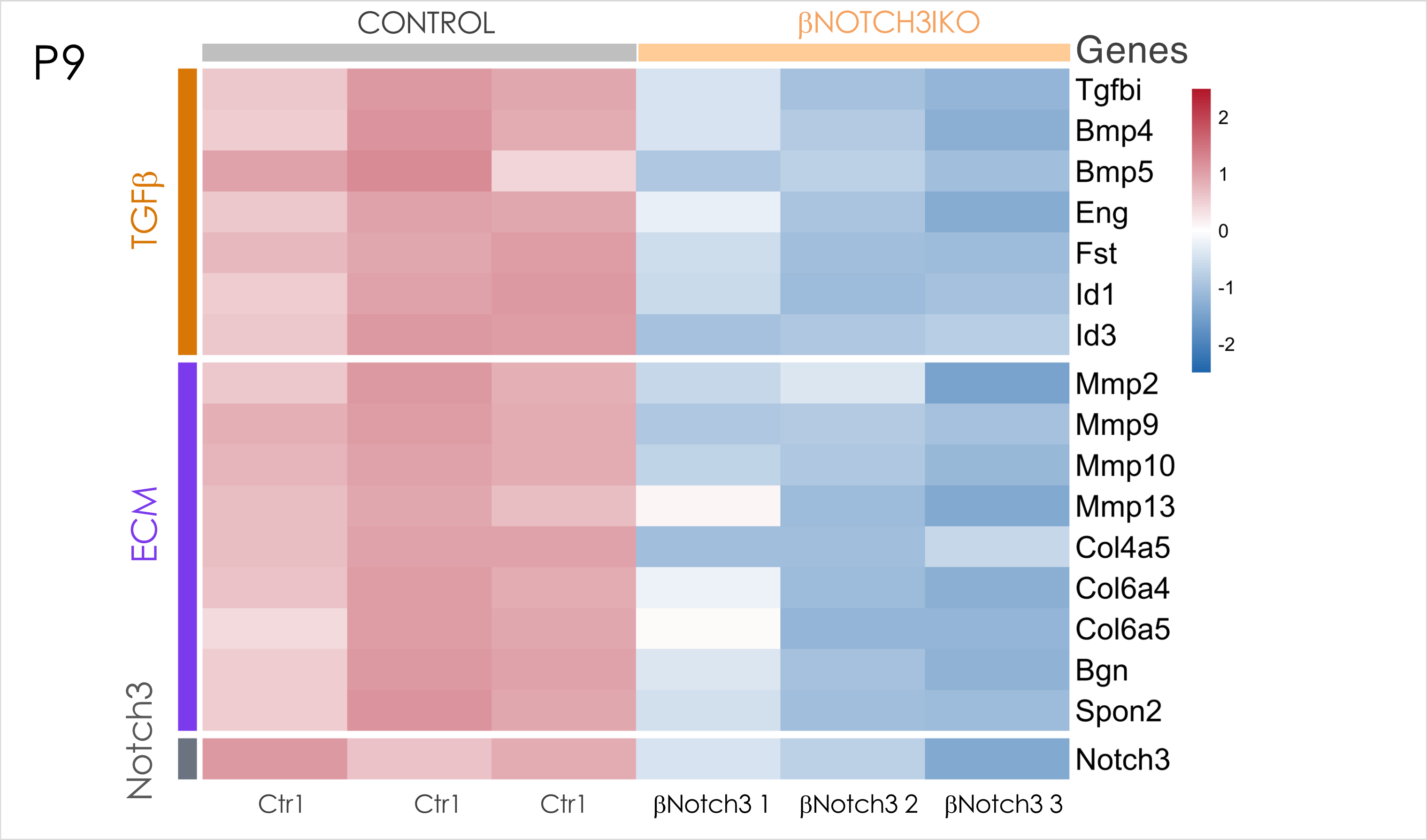


**C** GSEA reveals Notch3-dependent pathways including TGFβ signaling

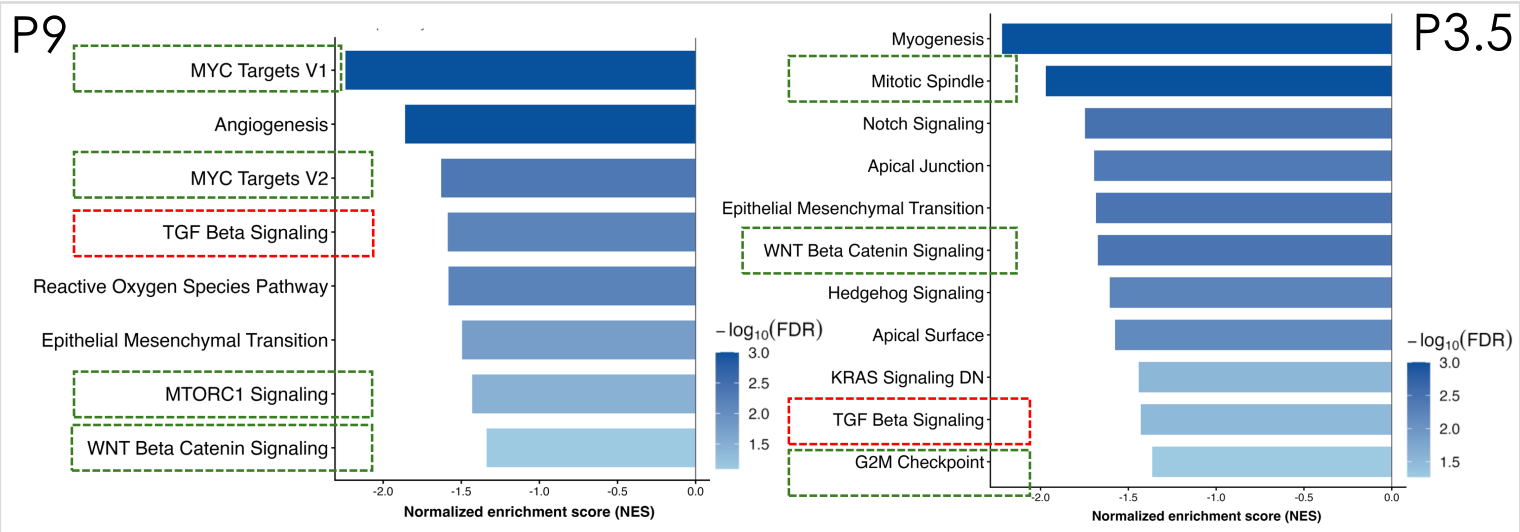
